# Circulating and brain-resident memory CD8+ T cells seed distinct bystander T_RM_-like populations in glioblastoma

**DOI:** 10.64898/2026.06.19.733403

**Authors:** Sierra A. Kleist, Tiffany Chen, Shawn C. Musial, Neva R. DiBlasi, Hanna N. Degefu, Sarah C. Berman, Myles A. Ford, Jordan F. Isaacs, Amanda Cruz Rivera, Arielle J. Sclar, Christina V. Angeles, Chun-Chieh Lin, Nathan E. Simmons, Linton T. Evans, Sladjana Skopelja-Gardner, Mary Jo Turk, Alexander G. J. Skorput, Sonia M. Leach, Pamela C. Rosato

**Author notes:** The authors declare no potential conflicts of interest.

## Abstract

Across cancers, tumor-infiltrating CD8+ T cells expressing the tissue-resident memory T cell (T_RM_) markers CD69 and CD103 are strongly associated with favorable clinical outcomes. However, a substantial fraction of these cells in human tumors are not tumor-specific, but instead recognize unrelated viral antigens. These virus-specific bystander T_RM_-like cells are prevalent in tumors and retain functional potential, raising interest in strategies that leverage pre-existing antiviral immunity for cancer immunotherapy. Yet their origins and differentiation states remain poorly defined, limiting both the interpretation of residency-based tumor-infiltrating lymphocyte (TIL) phenotyping and efforts to rationally harness these T_RM_-like cells. Here, using mouse models of GBM and melanoma, we demonstrate that resting circulating memory T cells trafficked into tumors via GPCR-dependent signaling and rapidly adopted a tissue-resident phenotype, independent of cognate antigen. Strikingly, in GBM, but not melanoma, pre-existing brain T_RM_ contributed substantially to the bystander TIL compartment and were the dominant source of CD69+/CD103+ bystander T cells, revealing a tumor- and tissue-specific origin for this subset. These findings were further supported by transcriptional analysis of T cell receptor clones present in both paired patient GBM tumor and peritumoral brain, which identified shared features with T_RM_-derived TILs in mouse GBM. Overall, this work provides new insight into tumor immunosurveillance, inform the interpretation of CD69+/CD103- and CD103+ TIL populations, and lay a foundation for immunotherapeutic strategies aimed at harnessing circulating and pre-existing virus-specific T_RM_ populations in tumors.

## INTRODUCTION

CD8+ T cells are key mediators of adaptive immunity, playing critical roles in both antiviral and antitumor responses. Following antigen clearance, a subset of these cells differentiates into long-lived memory populations with distinct migratory properties. Circulating memory T cells (T_CircM_) recirculate through lymphoid organs, peripheral blood, and sometimes tissues, while resident memory T cells (T_RM_) establish within lymphoid and non-lymphoid tissues and persist independently of continuous antigen exposure. T_RM_, often identified by expression of CD69 and CD103, are strategically positioned within tissues to mount more rapid local immune responses upon antigen re-encounter than their circulating counterparts (1,2). This capacity for local immune surveillance has generated considerable interest in the context of cancer. CD8+ tumor-infiltrating lymphocytes (TILs) with phenotypes resembling T_RM_ have been identified across numerous tumor types (3–5), and their presence has been associated with favorable clinical outcomes (2,3,6,7). However, not all TILs are tumor-reactive. Human tumors also contain non-tumor-specific, bystander CD8+ T cells that recognize microbial antigens unrelated to the tumor, including epitopes from common viruses such as Epstein-Barr virus (EBV), cytomegalovirus (CMV), and influenza A virus (8–10). Critically, these bystander T cells frequently display surface phenotypes overlapping with T_RM_ signatures commonly used to annotate antitumor immunity (8–10), potentially confounding the interpretation of TIL phenotyping data. Yet despite their prevalence, the origins, differentiation states, and residency properties of bystander CD8+ T cells within tumors remain poorly defined. This gap in knowledge is particularly consequential in glioblastoma (GBM), an immunologically ‘cold’ primary brain tumor characterized by limited TIL infiltration and poor responses to immunotherapy, where accurate interpretation of the immune landscape is needed.

Bystander T cells within the tumor microenvironment (TME) may nevertheless have important biological and therapeutic relevance. Previous studies have demonstrated that TME-associated bystander T cells remain unexhausted, suggesting they may serve as a latent reservoir of anti-tumor potential (9–11). For example, engagement of bystander T cells using a bispecific T cell engager antibody (BiTE), comprised of two single-chain variable fragments targeting CD3 and a tumor antigen, induced non-TCR-dependent T cell activation and reshaped the TME toward a more immunogenic state (12). In murine GBM models, antigen-activated non-tumor-specific T cells were capable of killing MHC-I-deficient tumor cells via NKG2D/NKG2D-ligand interactions (13). Moreover, direct intratumoral delivery of viral peptides, or the use of oncolytic viruses engineered to express tumor-irrelevant bystander T cell epitopes, have been shown to harness antiviral T cells to exert anti-tumor effects, although efficacy in GBM has been less consistent than in other malignancies (9,10,14,15). These findings suggest that virus-specific bystander T cells may influence tumor immunity even in the absence of cognate tumor antigen recognition, underscoring the importance of understanding their biology in tumors, including GBM.

T_RM_ form following both local infections, and distal infections/vaccinations in tissues throughout the body, including the brain (2,16,17). Because virus-specific memory T cells establish residency in tissues prior to malignancy, this provides a setting in which both pre-existing T_RM_ and recruited T_CircM_ may populate tumors and converge on a Trm-like phenotype, including expression of CD69 and CD103, complicating interpretation of their origin and migratory status. This is further compounded by the limitations of CD69 as a residency marker, as it is also rapidly upregulated by inflammatory signals and T cell activation, obscuring its use as a proxy for stable residency in an inflamed TME (18–21). This is particularly relevant in cancers that are difficult to detect early, such as GBM, where patients may harbor a tumor for months prior to diagnosis or definitive therapy (27), and may experience infections or vaccinations that reshape circulating and tissue-resident memory T cell pools over time, potentially seeding bystander populations that inflate or distort residency and activation readouts commonly used to infer tumor antigen recognition.

Several important questions therefore remain unresolved. First, the origins of bystander T_RM_-like TILs in tumors are incompletely understood; specifically, the relative contributions of pre-existing T_RM_ versus recruited T_CircM_ that subsequently adopt resident-like phenotypes within the TME have not been defined. Additionally, given that the CNS is known to impose unique constraints on lymphocyte trafficking, whether bystander T cell trafficking patterns differ between brain and peripheral tumors such as melanoma has not been established. It also remains unclear whether the CD69+/CD103-population in GBM represents stable tissue residency or activation-associated CD69 upregulation on newly recruited cells. Furthermore, because prior studies have relied heavily on mouse tumor models in which CD69+/CD103+ TILs are limited (22), it remains unknown whether CD103+ bystander TILs in tumors represent a compartment with distinct provenance and transcriptional properties relative to their CD69+/CD103-counterparts. Together, these gaps limit the interpretation of CD69+ and CD103+ TILs and highlight the need to define how non-tumor-specific memory CD8+ T cells contribute to the resident-like compartments within tumors.

Here, we used a reductionist murine system designed to separate tumor antigen recognition from T cell trafficking and residency-marker induction, allowing direct assessment of how distinct memory T cell compartments contribute to T_RM_-like CD8+ T cell states in GBM, using melanoma as a comparator to distinguish features unique to the CNS TME. Resting virus-specific T_CircM_ were found to traffic to GBM tumors in a GPCR-dependent manner, consistent with chemokine-mediated recruitment, and underwent rapid reprogramming towards a tissue-resident-like state. Moreover, pre-existing virus-specific T_RM_ residing in the healthy brain made up the majority of the CD103+ bystander T cell compartment in GBM, but not melanoma, and gave rise to a transcriptionally distinct resident-like population characterized by high granzyme expression. These findings provide a framework for interpreting CD69+/CD103- and CD103+ bystander populations within tumors and advance our understanding of brain tumor immune surveillance, laying the groundwork for future studies aimed at determining how pre-existing antiviral T cell immunity may be leveraged in GBM and other immunotherapy-resistant tumors.

## RESULTS

### Virus-specific memory T cells in tumors display a resident phenotype and increased abundance compared with healthy tissues

To better understand the phenotype and abundance of bystander, non-tumor-reactive TILs in a murine model of GBM, we generated mice with a trackable virus-specific memory T cell population. We adoptively transferred congenically marked (CD45.1+ or Thy1.1+) transgenic OT-I CD8+ T cells, specific for the SIINFEKL epitope of the model antigen ovalbumin (OVA), into naïve C57BL/6J mice followed by an intranasal infection with vesicular stomatitis virus expressing OVA (VSV_OVA_). VSV_OVA_ is an acute infection and establishes durable populations of circulating and resident memory T cells throughout the body within thirty days, including in the brain (23–25). These mice, herein referred to as “OT-I memory mice”, were then orthotopically implanted with a wild-type (OVA-negative) GBM cell line, GL261, known to harbor CD69+/CD103+ TIL population (10). A separate cohort of OT-I memory mice generated via i.v. VSVOVA infection received intradermal (i.d.) injections of the melanoma cell line B2905-M4 (M4) (51), to serve as a comparator, as we have found reproducible populations of CD69+/CD103+ bystander T cells in this tumor model in contrast to other flank tumors (22,26). Tumors were taken for flow cytometry at 15-17 days (GL261) or 21-28 days (M4), allowing sufficient time for tumor establishment and immune infiltration (Fig. 1a).

**Figure 1.**
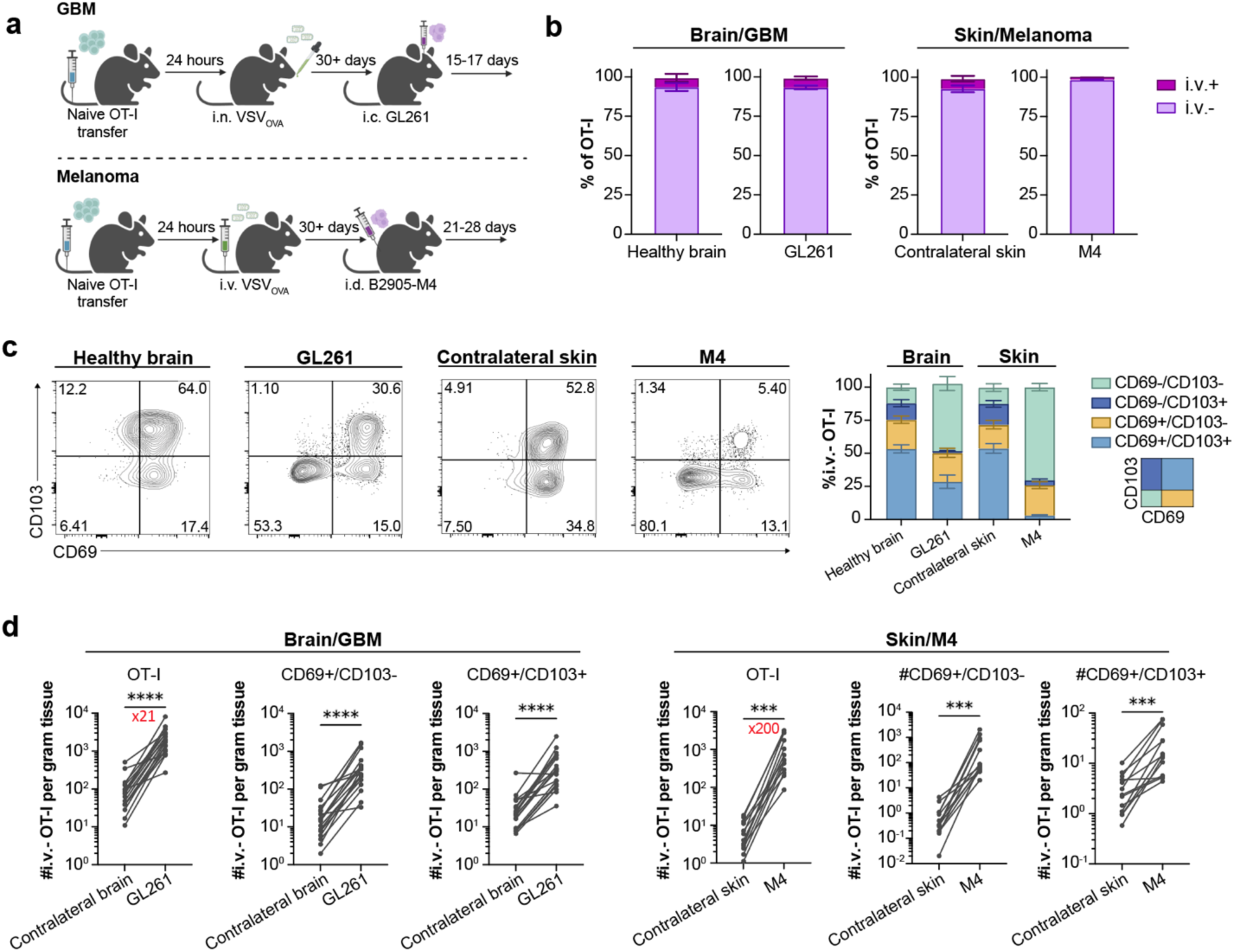
Virus-specific memory T cells in tumors display a resident phenotype and increased abundance compared with healthy tissues. a) Schematic of memory T cell generation and orthotopic tumor implantation. b) Bar graphs depicting i.v. antibody label of OT-I T cells isolated from brain, GL261 tumors, contralateral skin, and M4 melanoma tumors. c) Representative flow plots and quantification of CD69 and CD103 expression on OT-I T cells isolated from brain, GL261 tumors, contralateral skin, and M4 melanoma tumors. d) Absolute counts of total i.v.-negative OT-I, CD69+/CD103- and CD69+/CD103+ i.v.-negative OT-I T cells isolated from contralateral tissues and tumors. Fold change indicated in red text. GBM data in (b) and (c) are combined from two independent experiments; Healthy brain, *n* = 7; GL261, *n* = 19. M4 data in (b) and (c) are from three independent experiments; Contralateral skin, *n* = 13; M4, *n* = 13. Error bars represent the mean (±SEM) with *p* values determined by an unpaired t test or Mann Whitney test for non-normally distributed data. \*\*\**p* < 0.001, \*\*\*\**p* < 0.0001.

To distinguish between tumor parenchyma- and vascular-associated T cells, mice were injected i.v. with an anti-CD8α antibody three minutes prior to euthanasia to selectively label cells within the blood. In both tumor models, more than 90% of the OT-I T cells isolated from tumors were negative for the i.v. label, indicating that the majority of OT-I T cells were extravascular (Fig. 1b). We next compared the expression of residency-associated markers CD69 and CD103 on i.v.-negative OT-I cells in tumors and matched non-malignant tissues. To minimize potential confounding effects of tumor-associated inflammation on T_RM_ phenotype, GBM TILs were compared to healthy brain T_RM_ rather than to contralateral brain T_RM_, as GBM may induce CNS-wide changes (27). Contralateral skin served as an internal control in melanoma studies. Both GL261 and M4 tumors contained CD69+/CD103- and CD69+/CD103+ bystander TIL subsets, mirroring the two major T_RM_ populations found in non-malignant brain and skin (Fig. 1c), as well as in human tumors (9,10). However, both tumors also contained a substantial i.v.-negative CD69- /CD103-OT-I population, which was largely absent from healthy tissues (Fig. 1c). Quantification of i.v.-negative OT-I T cells revealed a 21-fold increase in GL261 tumors relative to contralateral brain, and a 200-fold increase in M4 tumors relative to contralateral skin (Fig. 1d). Both resident-like CD69+/CD103- and CD69+/CD103+ OT-I T cell subsets were also substantially enriched in tumors relative to non-malignant tissue (Fig. 1d). These data demonstrate that virus-specific TILs comprise both T_RM_-like subsets resembling those in corresponding non-malignant tissues, as well as a distinct i.v.-negative CD69-/CD103-population preferentially enriched in the TME.

### Resting virus-specific circulating memory T cells traffic to tumors and upregulate CD69

The significant increase in the number of CD69+ bystander TIL relative to contralateral tissue (Fig. 1d) suggests either recruitment from circulation with subsequent CD69 upregulation upon tumor entry, or incorporation of pre-existing CD69+ T_RM_ from the surrounding healthy tissue and local expansion within the tumor. Since circulating bystander T cell recruitment has been reported in extracranial tumors (11,22,28), we tested whether infiltrating virus-specific T_CircM_ contribute to CD69+ TIL populations in brain tumors. To generate mice with an intact T_CircM_ compartment, but lacking pre-existing T_RM_, we isolated circulating splenic CD8+ T cells from OT-I memory mice and adoptively transferred them to infection-naïve mice (Fig. 2a). Of note, T cells were isolated >30 days post-infection, a time when the virus is cleared and memory T cells are in a “resting” state. Transferred T_CircM_ were confirmed negative for expression of CD69 and CD103 (Fig. 2b). Within 24 hours of T_CircM_ transfer, mice were orthotopically injected with GL261 or M4 tumor cells and infiltrating T cells were assessed following tumor establishment (Fig. 2a).

**Figure 2.**
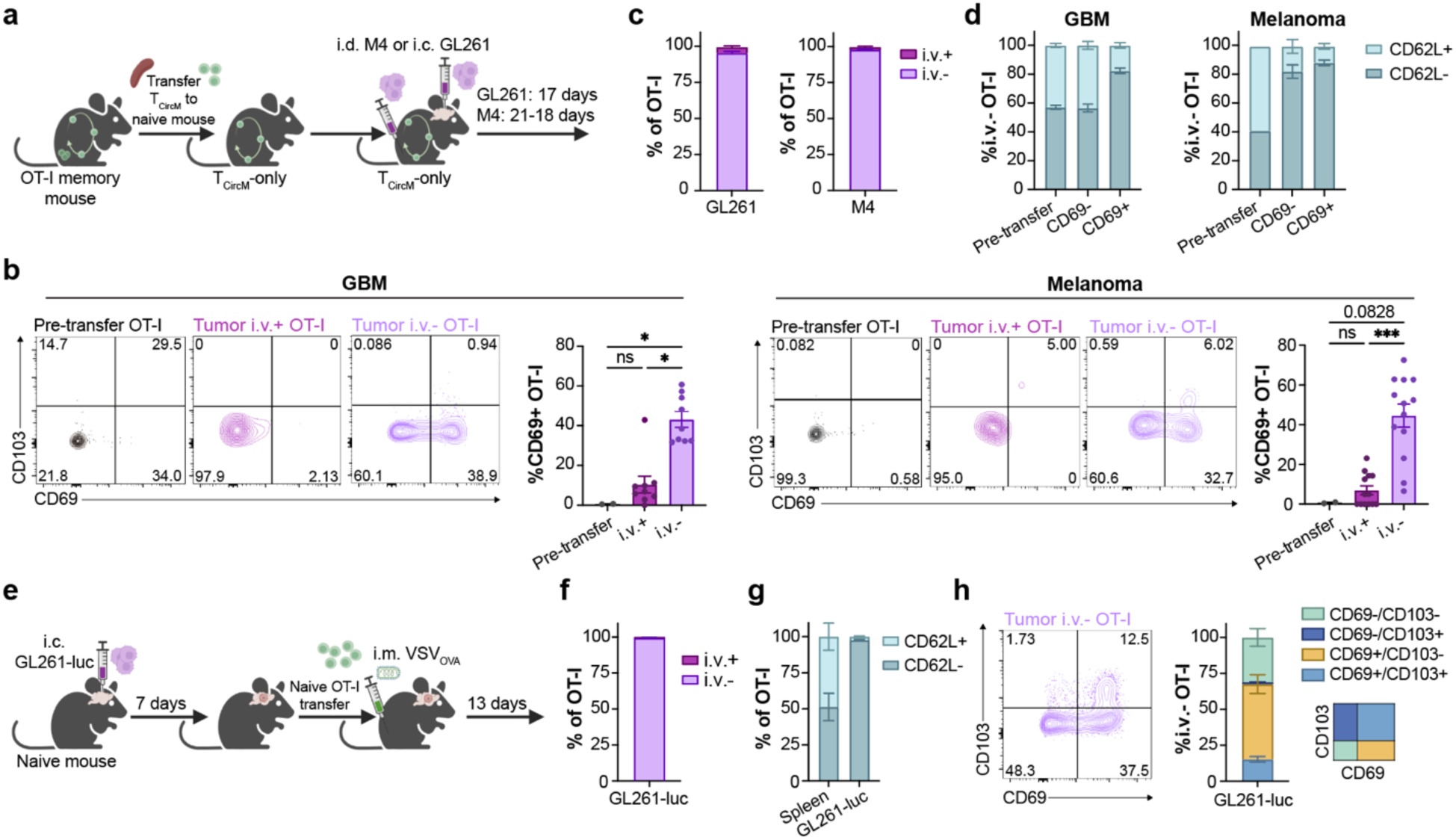
Resting virus-specific circulating memory T cells traffic to tumors and upregulate CD69. **a)** Schematic of T_CircM_ transfer to infection-naïve mice, followed by orthotopic tumor implantation. **b)** Representative flow plots and quantification of CD69 and CD103 expression on T_CircM_ before transfer (spleen) and following transfer (tumor) in the i.v.-label-positive and i.v.-label-negative populations. **c)** Bar graphs depicting i.v. antibody label of OT-I T cells isolated from tumors. **d)** Bar graphs showing distribution of CD62L- and CD62L+ subsets within transferred T_CircM_ prior to adoptive transfer (spleen) and after isolation from tumors. **e)** Schematic of intracranial GL261-luc injection followed by OT-I T cell transfer and intramuscular infection with VSV_OVA_. **f)** Bar graph depicting i.v. antibody label of OT-I T cells isolated from GL261-luc tumors 13 days post-infection. **g)** Bar graphs showing distribution of CD62L- and CD62L+ subsets within transferred T_CircM_ isolated from the spleen and from tumors 13 days post-infection. **h)** Representative flow plot and quantification of CD69 and CD103 expression on tumor-infiltrating OT-I T cells 13 days post-intramuscular infection. Data in (a) – (d) are combined from two independent experiments per tumor model; Pre-transfer, *n* = 2; Tumor OT-I, *n* = 9 (GL261) and *n*=13 (M4). Data in (e) – (h) are from one experiment; Tumor OT-I, *n* = 7. Error bars represent the mean ±SEM with *p* values determined by two-way ANOVA. \**p* < 0.05, \*\**p* < 0.01, \*\*\**p* < 0.001. ns, not significant.

Transferred OT-I T cells were readily recovered from GL261 brain tumors, with the majority being i.v.-negative, indicating that resting T_CircM_ can traffic to tumors and extravasate into the tumor parenchyma (Fig. 2b,c). Transferred OT-I T cells were also detected in flank M4 melanoma tumors (Fig. 2b,c), consistent with prior reports that resting bystander T_CircM_ infiltrate peripheral tumors (11,22). While OT-I T cells from the tumor vasculature (i.v.-positive) lacked expression of both CD69 and CD103, i.v.-negative OT-I T cells upregulated CD69 in both the GL261 and M4 models suggesting that T_CircM_ must exit the vasculature to upregulate this residency-associated markers (Fig. 2b). However, CD103 co-upregulation differed markedly between models, with GL261 tumors showing minimal CD103 expression (1.55% ± 0.26%), while M4 tumors showed low but detectable CD103 upregulation (5.24% ± 1.41%), a difference that was statistically significant between the two tumor models (*p* = 0.0176). While a mix of CD62L+ and CD62L-OT-I were recovered from the tumor, CD69+ cells were predominantly CD62L-in both models, consistent with a resident-memory like phenotype (Fig. 2d). Collectively, these findings demonstrate that resting bystander T_CircM_ can infiltrate brain and melanoma tumors and acquire a T_RM_-like phenotype upon tissue entry, despite lack of cognate antigen engagement.

We next asked whether recently activated virus-specific circulating CD8+ T cells can infiltrate established brain tumors and acquire a T_RM_-like phenotype, modeling a scenario in which tumor-bearing patients develop an infection or receive a vaccine. To model this in mice, we established intracranial GL261 tumors expressing firefly luciferase (GL261-luc). On day 7, following visualization of established tumors by IVIS imaging (Supplemental Fig. 1a,b), we adoptively transferred naïve OT-I T cells to the tumor-bearing mice followed by intramuscular or intranasal VSV_OVA_ inoculation, modeling a vaccination or infection, respectively (Fig. 2e and Supplemental Fig. 1a). At 13 days post-inoculation, the majority of tumor-derived OT-I T cells were CD62L- and i.v.-negative (Fig. 2f,g and Supplemental Fig. 1c,d). Additionally, a subset of infiltrating OT-I T cells expressed CD69, with some CD69+ cells co-expressing CD103 (Fig. 2h and Supplemental Fig. 1e), indicating that virus-specific effector T cells can also enter established brain tumors and acquire a resident-like phenotype.

### Recruitment of resting T_CircM_ to brain tumors is mediated by GPCR-signaling

To test if chemokine signaling drives the recruitment of resting bystander T_CircM_ to brain tumors, we performed a multiplex Luminex assay to quantify chemokines in GL261 tumors versus brain tissue from non-tumor bearing mice, where T_CircM_ trafficking is minimal. Numerous T cell recruiting chemokines, including CCL3, CCL4, CCL5, CXCL9, and CXCL10, were significantly enriched in brain tumors compared to healthy tissue (Fig. 3a). Chemokine receptors are part of the G protein-coupled receptor (GPCR) family. Therefore, to test if T_CircM_ trafficking to GBM tumors depends on chemokine receptor signaling, we cultured CD8-enriched splenocytes from OT-I memory mice with pertussis toxin (PTx) to block GPCR-signaling prior to adoptive transfer into mice bearing GL261 tumors (Fig. 3b). Twenty-four hours after i.v. transfer, PTx-treated T_CircM_ were absent from tumors, whereas untreated T_CircM_ robustly accumulated (Fig. 3c,d). Comparable frequencies of both PTx- and untreated-OT-I donor cells were recovered from the spleen, indicating that the reduced tumor trafficking was not due to a survival defect in PTx-treated T cells (Fig. 3c,d). Altogether, these data demonstrate that resting bystander T_CircM_ traffic to brain tumors in a GPCR-dependent manner, consistent with chemokine-mediated recruitment.

**Figure 3.**
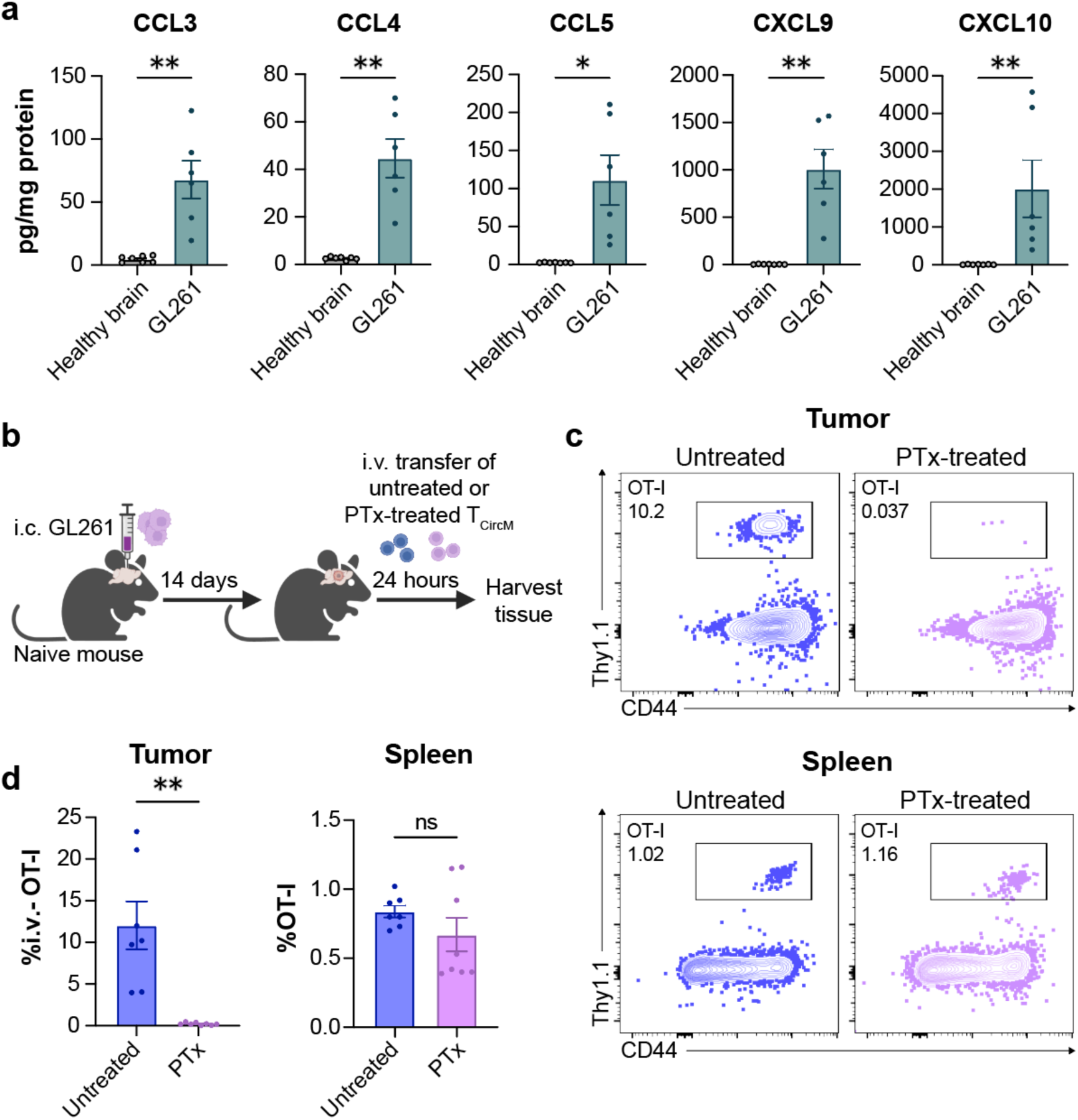
Recruitment of resting T_CircM_ to brain tumors is mediated by GPCR-signaling. **a)** Relative abundance of chemokines in GL261 brain tumors compared to healthy brain. **b)** Schematic of control or PTx-treated T_CircM_ transfer to GL261 tumor-bearing mice. **c)** Representative flow plots of untreated and PTx-treated CD44+ Thy1.1+ OT-I T cells isolated from tumor (top) and spleen (bottom), quantified in **(d).** Data in (a) are from two independent experiments; Healthy brain, *n* = 7; GL261, *n* = 6. Data in (b) – (d) are combined from two independent experiments; Untreated, *n* = 7; PTx treated, *n* = 8. Error bars represent the mean ±SEM with *p* values determined by an unpaired *t* test or Mann Whitney test for non-normally distributed data. \**p* < 0.05, \*\**p* < 0.01. ns, not significant.

### The tumor microenvironment rapidly alters the phenotype of bystander tumor-infiltrating T cells

CD69 is a canonical marker of tissue residency, but it is also upregulated upon T cell activation. Accordingly, CD69 expression on bystander T cells in tumors is challenging to interpret, as it may reflect inflammatory cytokine-driven activation or adoption of a tissue residency program. To distinguish these possibilities, we tracked the kinetics of CD69 upregulation together with described markers of cytokine-induced activation (granzyme B, CD25, Ki-67, NKG2D, IFNγ) (20,21,29) and inhibitory markers (PD-1, CD39), which are associated with T cells in tumors but are also highly expressed by functional brain T_RM_ (30–34). We used congenic markers to distinguish between transferred OT-I T cells, transferred endogenous CD8+ T cells, and host endogenous CD8+ T cells. Naïve CD45.1+/Thy1.1+ OT-I T cells were transferred into wild-type C57Bl/6J recipients followed by VSV_OVA_ infection. At 45+ days post-infection, CD8+ T cells were isolated from spleens of the OT-I memory mice and transferred i.v. to naïve CD45.1+/Thy1.1-mice bearing GL261 tumors. Tumors were then harvested at 24, 48, and 72 hours post-T_CircM_ transfer (Fig. 4a, Supplemental Fig. 2a).

**Figure 4.**
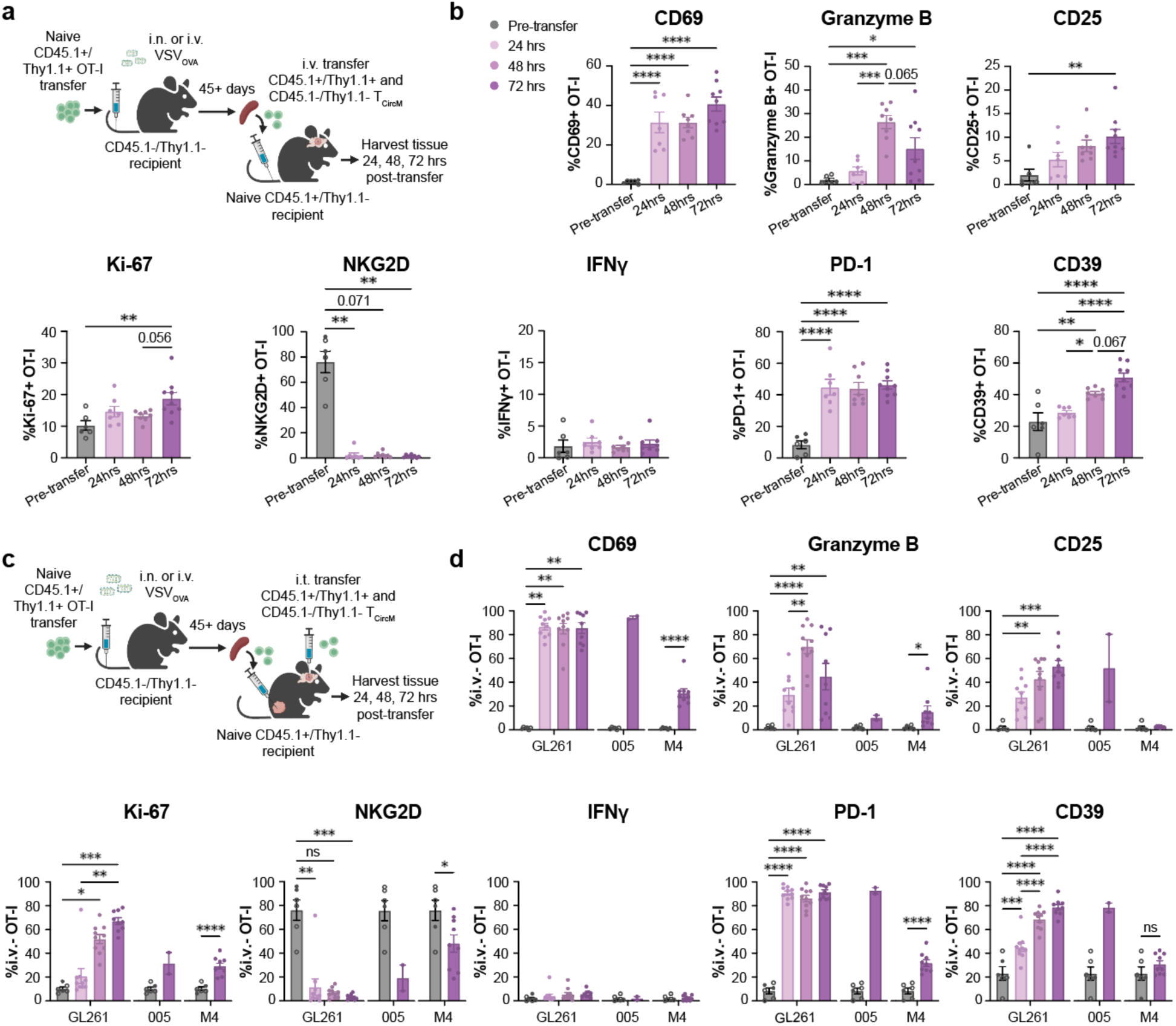
The tumor microenvironment rapidly alters the phenotype of bystander tumor-infiltrating T cells. **a)** Schematic of i.v. transfer of T_CircM_ to GL261 tumor-bearing mice. **b)** Expression of markers associated with residency and cytokine-mediated activation on OT-I T cells before (spleen) and from the tumor 24, 48, and 72 hours after i.v. transfer. **c)** Schematic of intratumoral transfer to mice bearing GL261 or 005 brain tumors, or M4 flank tumors. **d)** Frequency of OT-I expressing markers associated with residency and cytokine-mediated activation out of total OT-I T cells before (spleen) and from the tumor 24, 48, and 72 hours after i.t. transfer. Data in (b) are combined from two independent experiments; Pre-transfer, *n* = 6; 24hrs, *n* = 7; 48hrs, *n* = 8; 72hrs, *n* = 9. GBM data in (d) are combined from two independent experiments; Pre-transfer, *n* = 6; 24hrs, *n* = 10; 48hrs, *n* = 10; 72hrs, *n* = 9. 005 data in (d) are from one experiment; Pre-transfer, *n* = 6; 72hrs, *n* = 2. M4 data in (d) are from two independent experiments; Pre-transfer, *n* = 6; 72hrs, *n* = 9. Pre-transfer phenotypes of OT-I splenocytes (pooled from all experiments) were obtained from unmanipulated OT-I memory mouse splenocytes. Closed dots indicate cells used in the experiment shown; open circles represent cells from experiments with a different endpoint. Error bars represent the mean ±SEM with *p* values for GBM data determined by one-way ANOVA or Kruskal-Wallis test for non-normally distributed data and *p* values for 005 and M4 data determined with an unpaired *t* test or Mann Whitney test for non-normally distributed data. \**p* < 0.05, \*\**p* < 0.01, \*\*\**p* < 0.001, \*\*\*\**p* < 0.0001. ns, not significant.

Within 24 hours, a third of tumor-infiltrating T_CircM_ upregulated CD69, with expression maintained through 72 hours, alongside similar elevation of granzyme B, CD25, and Ki-67 (Fig. 4b, Supplemental Fig. 2b), consistent with the kinetics of cytokine-mediated activation (35,36). However, NKG2D, a receptor classically upregulated upon cytokine-mediated activation (21), was significantly downregulated within 24 hours of tumor entry. Moreover, IFNγ, which is typically expressed upon cytokine-mediated activation (21), was not detected, arguing against classical cytokine-driven activation (Fig. 4b, Supplemental Fig. 2b). PD-1 and CD39 expression also increased over 72 hours (Fig. 4b, Supplemental Fig. 2b). Taken together, the upregulation of CD69, granzyme B, PD-1, and CD39, combined with the downregulation of NKG2D and absence of IFNγ, is consistent with the acquisition of a residency program rather than classical cytokine-driven activation.

As i.v. delivered T_CircM_ may be continuously trafficking from the circulation to the tumor, newly arrived cells may not have been exposed to the TME for the full duration captured by each timepoint. To control for time of cell entry, we directly injected T_CircM_ intratumorally (i.t.) into GL261 and observed similar, but enhanced, upregulation of CD69, granzyme B, CD25, Ki-67, PD-1, and CD39, and loss of NKG2D in both OT-I T cells (Fig. 4c,d), and co-transferred endogenous memory (CD44^hi^) CD8+ T cells (Supplemental Fig. 2c,d). These results were reproduced when T_CircM_ were co-injected with an anti-MHC-I blocking antibody, supporting TCR-independent signaling, as expected in the absence of cognate antigen (Supplemental Fig. 3a,b). To determine if these findings were consistent across tumor types, we injected T_CircM_ into 005 tumors, a stem cell-derived invasive GBM model (37,38), and M4 flank tumors, and assessed OT-I T cells at 72 hours post-injection (Fig. 4c). In the 005 GBM tumors, we saw similar upregulation of CD69, CD25, PD-1, and CD39, and downregulation of NKG2D, but no increase in granzyme B, Ki-67, or IFNγ expression (Fig. 4d). In the M4 flank tumors, we also observed similar, but less robust, shifts in phenotype (Fig. 4d). Notably, CD25 was not upregulated on T_CircM_ in M4 tumors, and NKG2D was not reduced as robustly as was observed in the brain tumor models (Fig. 4d). The pronounced shift toward a T_RM_-like phenotype in the GL261 and 005 brain tumor models, compared with a similar but less robust shift in the M4 melanoma model, suggests that the brain TME may be particularly conducive to driving T_RM_-like differentiation of infiltrating T cells. Overall, these data indicate that the TME rapidly remodels infiltrating bystander T_CircM_ toward a T_RM_-like phenotype, in a TCR-independent fashion.

Type I interferons, IL-15, and IL-18, are well established to induce CD69 upregulation on memory T cells independently of TCR signaling. To test if these cytokines contributed to CD69 induction in our model, we neutralized IL-18, interferon alpha receptor (IFNAR), CD122 (a subunit of the IL-15 and IL-2 receptor), or a combination of IFNAR and CD122 with antibodies during i.t. transfer of T_CircM_ (Supplemental Fig. 4a). None of these treatments prevented CD69 upregulation or broader phenotypic shifts in transferred T_CircM_, mirroring the pattern observed previously (Supplemental Fig. 4b, Fig. 4). These data suggest the brain TME rapidly imposes a T_RM_-like program on infiltrating bystander T cells independently of IFNAR, CD122, and IL-18 signaling.

### Pre-existing brain T_RM_ seed the CD103+ bystander T cell compartment in brain tumors

Despite the prominent CD103+ bystander T cell population identified in GBM tumors (Fig. 1c), T_CircM_-derived OT-I T cells infiltrating GBM virtually lacked CD103 expression (Fig. 2b), suggesting an alternative origin for the CD103+ compartment. Prior parabiosis studies in murine models of breast cancer and melanoma reported that virus-specific TILs were derived primarily from circulation rather than from pre-existing T_RM_ in the corresponding healthy tissue (22). These models, however, lacked a prominent CD103+ TIL population, leaving open whether pre-existing T_RM_ contribute to the CD69+/CD103+ compartment in tumors that do harbor this subset. To test this, Thy1.1+ OT-I memory mice were treated with low-dose anti-Thy1.1 antibody to selectively deplete circulating OT-I T cells while sparing tissue-resident populations (Fig. 5a). Depletion was validated by the absence of OT-I T cells in the blood but retention in tissues (5b-d). Undepleted mice (T_RM_+T_CircM_) and T_CircM_-depleted mice were then orthotopically challenged with GL261 or M4 tumors. In GL261 tumors, the percent and total number of OT-I T cells infiltrating GL261 tumors was reduced by half in mice that received Thy1.1-depletion (Fig. 5e). In T_CircM_-depleted tumors, the frequency of CD103-expressing OT-I T cells was significantly increased, with a corresponding reduction in the CD69-/CD103-subset compared to undepleted control mice (Fig. 5f). Accordingly, the frequency and total number of CD69-/CD103-OT-I T cells was significantly reduced in T_CircM_-depleted mice, yet the number of CD69+/CD103+ OT-I T cells recovered from tumors remained unchanged between groups (Fig. 5f). In a complementary approach, we treated OT-I memory mice with FTY720, a sphingosine-1-phosphate (S1P) receptor modulator that blocks egress of T cells from lymph nodes, prior to GL261 implantation (Supplementary Fig. 5a). Consistent with depletion studies, FTY720 treatment significantly increased the frequency of CD103+ GL261-infiltrating OT-I T cells and reduced the CD69-/CD103-population, while the total number of CD69+/CD103+ OT-I T cells remained unchanged (Supplementary Fig. 5b-e). Together, these data indicate that the majority of the CD103+ OT-I T cells found in brain tumors derive from pre-existing brain T_RM_.

**Figure 5.**
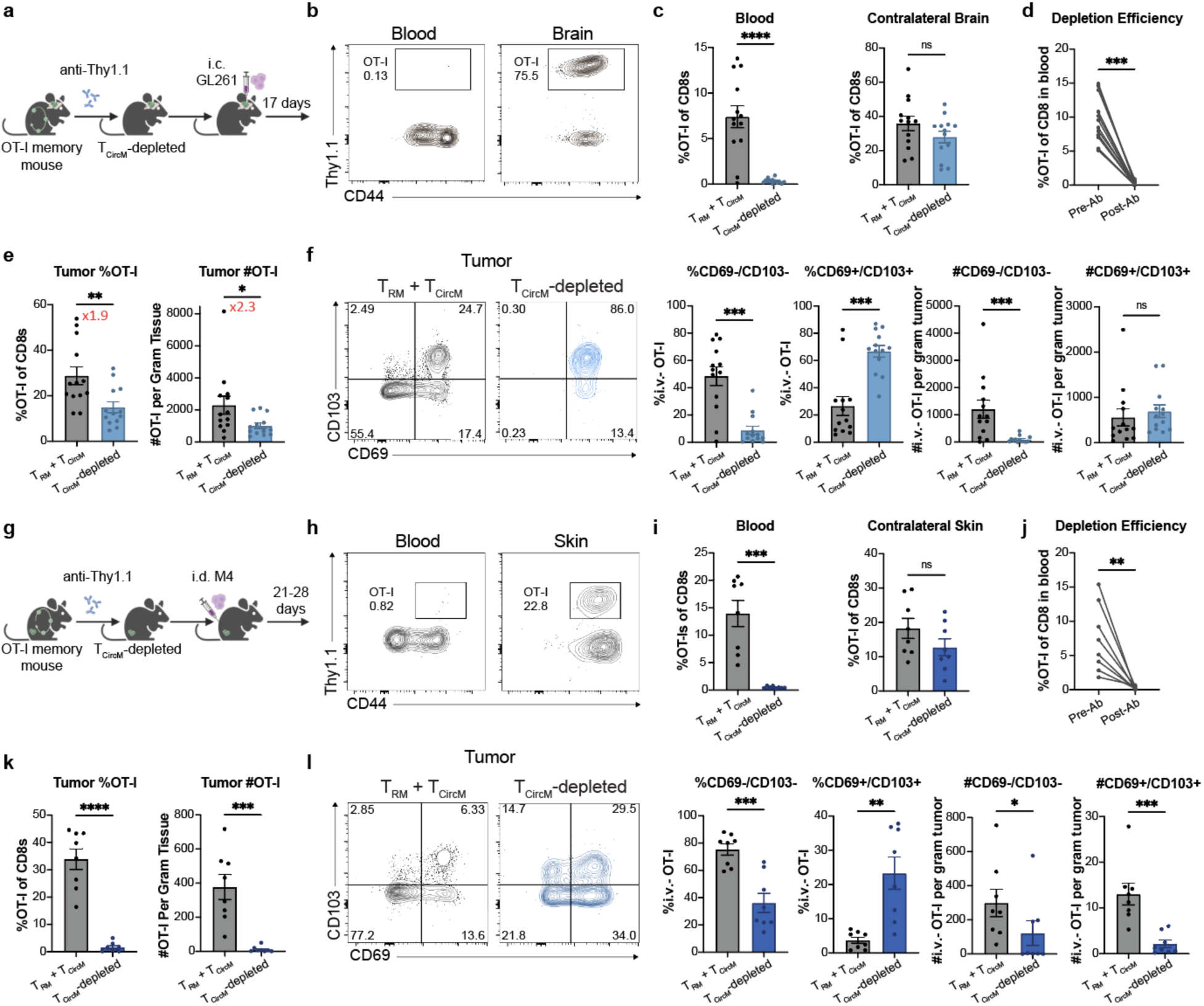
Pre-existing brain T_RM_ seed the CD103+ bystander T cell compartment in brain tumors. **a,g)** Schematic of T_CircM_-depletion and tumor implantation. **b, h)** Representative flow plots depicting Thy1.1+ CD44^hi^ OT-I T cells in blood and tissue from mice receiving αThy1.1 depleting antibody or undepleted controls (T_RM_+ T_CircM_), quantified in **(c, i)**. **d, j)** OT-I frequencies in the blood before and after antibody depletion. **e, k)** Frequency and total number of OT-I T cells isolated from tumors. Fold change indicated in red text. **f, l)** Representative flow plots and quantification of CD69 and CD103 expression on OT-I T cells in tumors from undepleted mice (T_CircM_ + T_RM_) and T_CircM_-depleted mice. GBM data are combined from three independent experiments. T_RM_ + T_CircM_, *n* = 13; T_CircM_-depleted, *n* = 13. M4 data are combined from two independent experiments; T_RM_ + T_CircM_, *n* = 8; T_CircM_-depleted, *n* = 8. Error bars represent the mean ±SEM with *p* values determined by unpaired *t* test or Mann Whitney test for non-normally distributed data. *p* values in (d) and (j) determined with paired *t* test. \**p* < 0.05, \*\**p* < 0.01, \*\*\**p* < 0.001, \*\*\*\**p* < 0.0001. ns, not significant.

In stark contrast to the brain tumor model, T_CircM_ depletion dramatically reduced OT-I T cell recovery from M4 melanoma tumors (Fig. 5g-k). However, among the cells recovered, similar shifts in CD69 and CD103 expression were observed, with a decreased frequency of CD69- /CD103-OT-I T cells and an increased frequency of CD103+ cells, consistent with the GL261 model (Fig. 5l). Thus, while depletion of T_CircM_ enriches for CD103 expression among melanoma OT-I TILs, pre-existing skin T_RM_ are not the dominant source of bystander OT-I T cells in M4 tumors. Overall, these findings suggest that the contribution of pre-existing T_RM_ to TILs varies by tumor type and anatomic site. Nevertheless, pre-existing T_RM_ generally bias the antiviral TIL compartment toward CD103 expression and, in GBM, account for a substantial fraction of total bystander TILs.

### Brain tumor T_CircM_-derived TILs acquire a resident-like transcriptional program, whereas TILs from pre-existing T_RM_ preserve a distinct resident signature

To define how T_CircM_ are transcriptionally remodeled upon entry into GL261 brain tumors, and to determine whether these cells converge on the same transcriptional states as TILs derived from pre-existing brain T_RM_, single-cell RNA-sequencing (scRNA-seq) was performed on sorted OT-I TILs from two complementary conditions: mice that received an i.v. T_CircM_ transfer (T_CircM_-only) and mice depleted of T_CircM_ (T_RM_-only). OT-I T cells from the spleens of tumor-bearing OT-I memory mice were also profiled as a circulating comparator. Cells from each group were multiplexed using oligo-hashtag barcodes prior to pooling to enable group assignment.

Unsupervised clustering across the three experimental groups identified eight transcriptionally distinct clusters (C0–C7), visualized by UMAP (Fig. 6a). Overlay of experimental group identity and quantification of cluster composition showed that OT-I TILs from both T_CircM_-only and T_RM_-only conditions were largely transcriptionally distinct from the circulating spleen clusters, yet overlapped extensively with one another (Fig. 6b-d), indicating that entry into the GBM microenvironment is sufficient to reprogram T_CircM_ toward a resident-like transcriptional state. As expected, spleen-derived OT-I T cells were enriched in clusters expressing canonical circulating genes; C5 and C6 expressed effector memory (T_EM_)-associated transcripts, including *Cx3cr1* and *Klrg1*, while cells from C7 expressed a central memory (T_CM_) program characterized by high *Ccr7, Sell* (CD62L), *S1pr1*, *Il7r*, *Slamf6*, and *Tcf-7* (TCF-1) (Fig. 6a,e).

**Figure 6.**
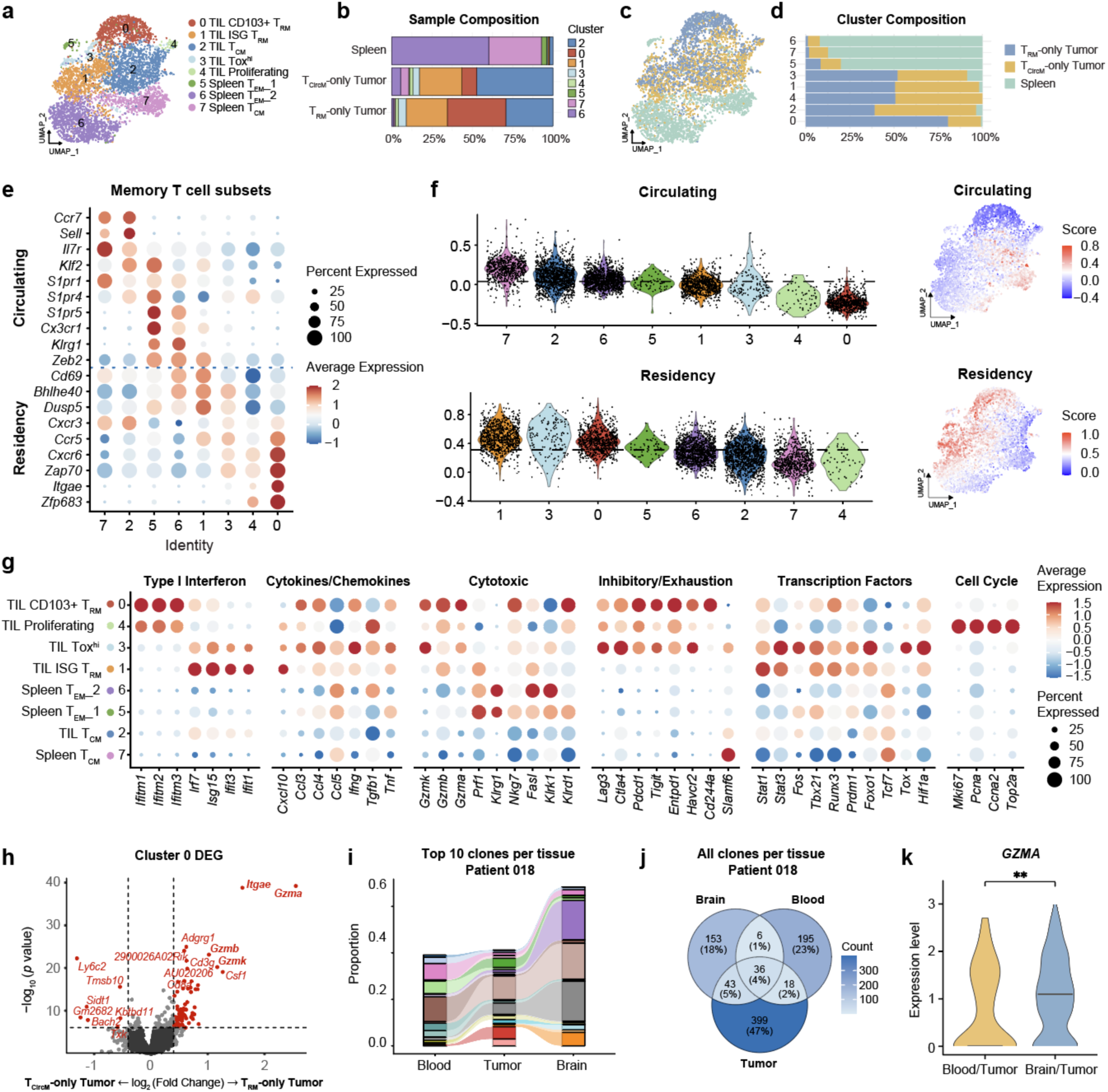
Brain tumor T_CircM_-derived TILs acquire a resident-like transcriptional program, whereas TILs from pre-existing T_RM_ preserve a distinct resident signature. scRNA-seq analysis of sorted virus-specific memory OT-I T cells from T_RM_-only tumors (T_CircM_-depleted), T_CircM_-only tumors, and spleen 15 days after tumor challenge. **a)** UMAP of transcriptional states annotated by cluster name. **b)** Relative frequencies of OT-I T cell clusters in each experimental group/tissue. **c)** UMAP of OT-I T cells annotated by experimental group/tissue. **d)** Relative frequencies of each experimental group/tissue in each cluster. **e)** Dot plot of select circulating and resident memory T cell transcripts across clusters. **f)** Violin plots (left) and feature plots (right) depicting circulating and residency module scores across clusters. Module scores generated from Gavil et al. (22). **g)** Dot plot of select cluster-defining genes. **h)** Volcano plot of DEGs between Cluster 0 OT-I T cells from T_CircM_-only v.s. T_RM_-only tumors. **i)** Alluvial plots depicting overlap of the top 10 CD8+ T cell clonotypes across Patient 018 blood, high grade glioma tumor, and peritumoral brain, with flow width proportional to clone frequency. **j)** Venn diagram showing all shared TCR clonotypes across Patient 018 tissues. **k)** Violin plots comparing *GZMA* expression in tumor-infiltrating CD8+ T cells belonging to blood-tumor shared clonotypes vs. peritumoral brain shared clonotypes. Data in (a – h) are from one independent experiment, with cells in each group pooled from *n* = 8-10 mice. High grade glioma patient data in (i-k) are from Patient #P018, *n* = 1. *p* value determined by Wilcoxon on SCT normalized data as used in Seurat FindMarkers(). \*\**p* < 0.01.

Among the tumor-enriched clusters (C0-C4), both circulating and tissue-resident transcriptional profiles were observed, supported by module scoring with published circulating and residency gene signatures (Fig. 6e,f) (22,39). C2 scored high for a circulating gene signature, consistent with elevated expression of *Ccr7*, *Sell*, and *Tcf7* and a T_CM_-like phenotype (Fig. 6e,f), mirroring the CD69-/CD62L+ tumor-infiltrating population identified by flow cytometry (Fig. 2). This cluster was enriched in the T_CircM_-only sample (Fig. 6b,d), suggesting that many of these cells are recent tumor entrants that have not yet acquired a residency program. However, C2 was also detected in the T_RM_-only sample, consistent with the hypothesis that pre-existing T_RM_ exhibit developmental plasticity within the inflammatory TME and may transition toward an ex-T_RM_ phenotype, as seen in other contexts (40,41).

The remaining tumor-enriched clusters, C0, C1, C3, and C4, expressed transcripts associated with tissue residency (Fig. 6e), though module scoring supported this classification only for C0, C1, and C3 (Fig. 6f). C4, by contrast, showed low scores for both residency- and circulating-associated modules and clustered primarily based on cell cycle gene expression, consistent with a small proliferating T_RM_-like subset (Fig. 6f,g). C3 corresponded to a resident but inhibited population, scoring highly for residency-associated modules while also expressing an exhaustion program, including *Lag-3*, *CTLA-4*, *Havcr2*, and the transcription factor *Tox* (Fig. 6f,g). C1 comprised equal proportions of T_CircM_-only and T_RM_-only cells and exhibited high expression of *CD69*, *Bhlhe40*, and *Dusp5* (Fig. 6e), as well as interferon-stimulated genes (*Ifit1*, *Ifit3*, *Isg15*, and *Irf7*) (Fig. 6g). Consistent with the CD69+/CD103-TIL population identified by flow cytometry (Fig. 1), C1 exhibited the highest residency-associated gene module score among all clusters (Fig. 6f), further supporting that T_CircM_ can acquire a resident-like transcriptional state upon tumor entry.

C0 was comprised predominantly of OT-I T cells from T_RM_-only tumors (∼75%), with the remainder derived from T_CircM_-only tumors (Fig. 6d). C0 exhibited a distinct transcriptional profile characterized by the highest expression of the T_RM_-associated genes *Itgae* (CD103), *Zfp683* (Hobit), *Zap70*, and *Cxcr6* (Fig. 6e), and the lowest circulating gene module score among all clusters (Fig. 6f). This cluster was additionally distinguished by high expression of interferon-inducible transmembrane genes (e.g. *Ifitm1-3*), cytotoxic effector molecules (*Gzma, Gzmb, Gzmk*), and inhibitory molecules (*Pdcd1*, *Tigit*, *Entpd1*, *Havcr2*, *Cd244a*) (Fig. 6g). Given that C0 was comprised of ∼75% OT-I T cells from T_RM_-only tumors, differential gene expression analysis was performed to determine whether these cells maintained a transcriptionally distinct identity relative to OT-I T cells from T_CircM_-only tumors within the same cluster. Cells from T_RM_-only tumors exhibited significantly higher expression of *Itgae*, *Gzma*, *Gzmb*, and *Gzmk* (Fig. 6h), indicating that pre-existing T_RM_ retain a distinct transcriptional identity within this cluster. Altogether, these data support a model in which T_CircM_ entering GBM can adopt a resident-like transcriptional state, while pre-existing brain T_RM_ give rise to a distinct resident compartment that is uniquely enriched for residency-associated genes and poised for cytotoxicity.

To assess whether distinct memory T cell compartments similarly give rise to transcriptionally distinct T_RM_-like cells human GBM, TCR sequencing was performed on CD8+ T cells isolated from paired blood, tumor, and peritumoral brain tissue from a patient with high grade glioma undergoing routine surgical resection. We used TCR clonotype sharing across anatomic compartments as a surrogate for potential relationships between circulating, tumor-infiltrating and brain CD8+ T cell populations. Specifically, clones shared only between blood and tumor were interpreted as putative circulating-derived tumor-infiltrating clones, whereas clones shared only between tumor and peritumoral brain were interpreted as putative brain-derived tumor-infiltrating clones. The majority of clones were shared across all three tissues; however, we identified clones shared exclusively between blood and tumor, and exclusively between tumor and peritumoral brain (Fig. 6i,j). Notably, within the tumor sample, T cells belonging to clonotypes shared exclusively between peritumoral brain and tumor expressed higher *GZMA*— a key defining gene of brain T_RM_-derived TILs in murine GBM (Fig. 6h)— relative to tumor T cells belonging to clonotypes exclusively shared between blood and tumor (Fig. 6k). Although antigen specificity remains unknown, and this analysis was limited to a single patient, these data provide initial evidence that the distinct origins of T_RM_-like populations identified in murine GBM may be conserved in human disease.

## DISCUSSION

In this study, we examined how distinct virus-specific memory T cell populations contribute to T_RM_-like cell states in murine models of GBM and melanoma. CD69+/CD103- and CD69+/CD103+ bystander T cells accumulated in both tumor models, initially suggesting that resting T_CircM_ traffic to tumors and acquire a residency-associated phenotype, as proposed previously (11,22,42). Consistent with these studies, we found that the CD69+/CD103-compartment was predominantly composed of T_CircM_-derived cells in both GBM and melanoma models. However, the murine tumor models in those prior studies largely lacked a CD103+ TIL population, a subset that is prominent in multiple human tumors, including GBM (9,10). Using GBM and melanoma models that do harbor CD103+ TILs, we found that the cellular origins of this compartment differed by anatomic site; In melanoma, T_CircM_ were the dominant source of T_RM_-like bystander T cells and were sufficient to generate a CD103+ compartment. In contrast, while resting T_CircM_ entering brain tumors acquired resident-like features, including expression of CD69, PD-1, and loss of CD62L, pre-existing T_RM_ from the healthy brain contributed a substantial proportion of TILs and preferentially gave rise to the CD103+ compartment. Together, these findings suggest that the brain may be a distinct tumor site in which pre-existing tissue-resident memory T cells, rather than T cell recruitment from the circulation alone, can substantially shape the bystander TIL compartment.

Building on these observations, we next investigated the mechanisms governing bystander T cell entry into brain tumors. We found that resting bystander T cells required GPCR-mediated signaling to infiltrate brain tumors, likely in response to the abundant chemokines present within the TME identified here and by others (43,44). Previous studies have shown that activated bystander T cells, which express heightened levels of trafficking molecules, infiltrate peripheral tumors through CXCR3- and CCR5-dependent signaling (28,45). In GBM, CCR5 may play a less prominent role, as prior studies did not observe a defect in CD8+ T cell infiltration in CCR5-/-mice bearing GL261 tumors (43). In contrast, the CXCR3 ligand CXCL10 appears to be important for recruitment of activated bystander T cells to brain tumors, as its neutralization in a melanoma brain metastasis model reduced CD8+ T cell infiltration (46), and increasing intratumoral CXCL10 enhanced CD8+ T cell accumulation in GBM models (47). Whether similar mechanisms govern the recruitment of *resting* bystander T_CircM_ to brain tumors remains unknown. In peripheral viral infection models, resting bystander T cells traffic to inflamed tissue in a CXCR3-dependent manner (35), and CXCR3 expression on resting T_CircM_ is required for trafficking to peripheral murine tumors (42). Although CXCR3 was not directly tested in this study, our finding that resting bystander T cells require GPCR-mediated signaling to infiltrate brain tumors, together with the above studies, suggests that CXCR3 may similarly contribute to bystander T_CircM_ trafficking to GBM.

Upon tumor entry, bystander T_CircM_ underwent rapid phenotypic remodeling. Within 24 hours of entry, tumor-infiltrating T_CircM_ upregulated of a set of surface markers, including CD69, CD25, Granzyme B, PD-1, and CD39, a pattern interpretable as either cytokine-mediated activation or initiation of a tissue residency program. Although the inflammatory milieu of the tumor could in principle trigger activation of newly recruited cells, analogous to cytokine-mediated activation during viral infections, classical markers of such activation, including NKG2D and IFNγ, were not detected. Furthermore, blockade of cytokine signals known to drive cytokine-mediated activation failed to prevent CD69 upregulation on tumor-infiltrating T cells, further supporting the adoption of a residency-associated rather than an activation phenotype. Consistent with this interpretation, a subset of tumor-infiltrating T_CircM_ was enriched for tissue-residency transcriptional signatures, revealing a previously unappreciated capacity of resting T_CircM_ to infiltrate and adopt a resident-like state within brain tumors. The factors driving this reprogramming remain undefined, but may include hypoxia, which has been shown to induce CD69 expression in cultured human PBMCs and mouse splenocytes (48,49).

While T_CircM_ gave rise primarily to the CD69+/CD103− compartment, the CD103+ population had a distinct origin. We found that pre-existing brain T_RM_ give rise to CD103-expressing T cells in brain tumors, identifying an unexplored source of TILs distinct from circulation. This contrasts with peripheral melanoma tumors, where very few cells from the skin T_RM_ pool infiltrated tumors, consistent with previous reports (22). Notably, intramuscular vaccination of mice bearing brain tumors was similarly sufficient to generate CD103+ TILs. These CD69+/CD103+ TILs maintained features distinct from their CD69+/CD103-counterparts, including enriched expression of residency-associated genes (*Zfp683* (Hobit), *Itgae* (CD103), *Zap70*, *Cxcr6*) and cytotoxicity-associated genes (*Gzma*, *Gzmb*, *Gzmk)*, suggesting this population is uniquely poised for cytotoxicity. Future studies are needed to elucidate the functional differences between these distinct CD69+/CD103- and CD69+/CD103+ T_RM_-like states.

These findings carry important implications for how we think about human brain immunity. As human brain tumors contain a substantial population of CD103+ T_RM_-like cells specific for common pathogens (9,10), our results offer a potential framework for interpreting their origin. If the observations reported here extend to humans, the virus-specific CD69+/CD103+ T_RM_-like cells found in GBM may reflect the T_RM_ landscape present in healthy human brain prior to malignancy, a possibility of particular interest given that the antigen specificity of T_RM_ in normal human brain remains poorly defined. In support of this concept, analysis of matched samples from an individual with high-grade glioma revealed that CD8+ TCR clones shared between tumor and peritumoral brain tissue were enriched for *GZMA*, a marker that distinguished the T_RM_-derived population in our murine dataset. In contrast, clones shared between tumor and blood lacked this signature, consistent with a distinct, circulation-derived origin. Although these observations are limited to a single patient and do not establish antigen specificity, they suggest that the cellular origins we defined in mice may be reflected in human brain tumors. Our study further raises the possibility that patients with brain tumors may acquire new bystander T_RM_-like cells within tumors following viral infection or vaccination, an important consideration as the field continues to define the role of bystander T cells in anti-tumor immunity.

Altogether, this study defines the origins of virus-specific T_RM_-like cells in murine brain tumors and provides new insight into the composition and interpretation of CD69+/CD103- and CD69+/CD103+ T cell populations within GBM. By clarifying the cellular dynamics that shape T cell residency in the brain TME, this work advances our understanding of immune surveillance in the CNS and may inform future immunotherapeutic strategies that seek to harness bystander T cell immunity in brain tumors.

## MATERIALS AND METHODS

### Human subjects

Institutional review board-approved written, informed consent was obtained from patients scheduled for routine open cranial resection of brain tumors as standard of care deemed by the treating neurosurgeon. All human studies were conducted in accordance with ethical regulation and preapproved by the Committee for the Protection of Human Subjects at Dartmouth-Hitchcock Medical Center (no. 02001404).

### Mice

C57Bl/6J mice were purchased from The Jackson Laboratory (Bar Harbor, ME) and were maintained at Dartmouth College under standard housing, husbandry, and dietary conditions according to the Institutional Animal Care and Use Committee (IACUC) and NIH guidelines. OT-I transgenic mice were originally purchased from The Jackson Laboratory, were fully backcrossed to C57BL/6J mice, and were maintained in our animal colony on a Thy1.1+ and/or CD45.1+ congenic background. Age- and sex-matched mice were used for all experiments. Mice used in experiments were 8-20 weeks of age. Both male and female mice were used in initial experiments. As no sex-dependent differences were observed, subsequent experiments were performed using only female mice. No sample exclusion criteria was applied. All experimental procedures performed were approved by and in accordance with the Institutional Animal Care and Use Committees at Dartmouth College.

### Adoptive transfers and infections

Immune memory mice were generated by adoptively transferring 10,000 or 50,000 Thy1.1+ or CD45.1+ CD8+ OT-I T cells from male or female mice into sex-matched naïve 6-8-week-old CD45.2+ C57Bl/6J recipients via tail vein. One day following transfer, mice were infected intranasally (i.n.) with 5×10⁴ PFU or intravenously (i.v.) with 1×10⁶ PFU of vesicular stomatitis virus expressing chicken ovalbumin (VSV_OVA_).

### Lymphocyte isolation and phenotyping

We used an intravascular staining method to distinguish cells present in the vasculature from cells present in the tissue parenchyma as previously described (50,51). Mice were injected with 3μg of biotin-conjugated anti-CD8α (clone 53-6.7, BioLegend 100703) via tail-vein. Three minutes after the injection, the mice were euthanized and tissues were harvested as described (51).

GL261 tumors, B2905-M4 tumors, contralateral brain, and contralateral skin were removed, cut into small pieces, and digested in Collagenase IV (Sigma) (0.5mg/ml for tumors and brain, 3mg/ml for skin) and DNAse I (5mg/ml) for 30 minutes at 37°C. All tissues were then mechanically disrupted using a GentleMacs dissociator (Miltenyi Biotec) and filtered through 70um mesh. Lymphocytes were then isolated using a 44/67% Percoll (Cytiva) density gradient and stained for 30 minutes at 4°C with antibodies to Thy1.1-BUV737 (clone HIS51, BD Biosciences 741774), Ki-67-eFluor 450 (clone SolA15, Thermo Fisher Scientific 48-5698-82), IFNγ-BV421 (clone XMG1.2, BioLegend 505829), CD103-BV510 (clone 2E7, BioLegend 121423), CD44-BV605 (clone IM7, BioLegend 103047), CD45.1-BV650 (clone A20, BioLegend 110735), CD44-BV785 (clone IM7, BioLegend 103059), CD39-PerCP-eFluor710 (clone 24DMS1, Thermo Fisher Scientific 46-0391-80), CD45.1-Alexa Fluor 488 (clone A20, BioLegend 110718), CD44-FITC (clone IM7, BioLegend 103006), Granzyme B-PE (clone GB11, Thermo Fisher Scientific 12-8899-41), Granzyme B-PE (clone GB11, BD Biosciences 561142), CD69-PE-Dazzle594 (clone H1.2F3, BioLegend 104535), PD-1-PE-Cy7 (clone 29F.1A12, BioLegend 135216), CD62L-PE-Cy7 (clone MEL-14, BioLegend 104418), NKG2D-APC (clone CX5, BioLegend 130212), CD8β-Alexa Fluor 647 (clone YTS156.7.7, BioLegend 126611), CD25-APC-Cy7 (clone PC61, BioLegend 102025), Streptavidin-BV605 (BioLegend 405229), Streptavidin-BV650 (BioLegend 405232), and viability dyes LIVE/DEAD Fixable Blue Dead Cell Stain Kit (Thermo Fisher Scientific L23105), Ghost Dye Red 780 (Cytek Biosciences Inc 13-0865-T100) for 30 minutes at 4°C for flow cytometry. Cells stained intracellularly were permeabilized using the Tonbo Biosciences Foxp3/Transcription Factor Staining Buffer Kit. Enumeration was done using Invitrogen CountBright™ Plus Absolute Counting Beads. Stained samples were acquired using a Cytek Aurora and analyzed with FlowJo software.

### Tumor models and treatment

GL261 cells were obtained from the Division of Cancer Treatment and Diagnosis (DCTD) Tumor Repository, National Cancer Institute (NCI) (Bethesda, MD). GL261-luc cells were shared by Dr. Tyler Curiel (Dartmouth), and B2905-M4 cells were shared by Dr. Mary Jo Turk (Dartmouth) (52), and 005 cells were shared by Dr. Susan Kaech (Allen Institute) (37,38). GL261 and GL261-luc cells were cultured in DMEM supplemented with 10% FBS and 1% penicillin/streptomycin (Corning). 005 glioma stem cells were cultured in DMEM/F12 supplemented with 2 mM L-glutamine (Corning), 1% N2 supplement (Gibco), 1% penicillin/streptomycin (Corning), recombinant human EGF (20 ng/mL; PeproTech), and recombinant human FGF-basic (20 ng/mL; PeproTech). B2905-M4 cells were cultured in RPMI supplemented with 10% FBS. All cell lines were cultured at 37°C in 5% CO_2_. Cells were dissociated with TrypLE^TM^ Express (ThermoFisher) for passaging.

To establish brain tumors, 100,000 GL261, 300,000 GL261-luc, or 300,000 005 cells were injected intracranially into mice. Briefly, a hand drill mounted on a stereotaxic frame was used to make a burr hole 1mm left and 1mm posterior of bregma, taking care not to damage the underlying dura mater. A Hamilton syringe with a 26-gauge needle was guided by the stereotaxic frame to the coordinates of the burr hole, and lowered 2mm deep into the cerebral hemisphere. 3μL of cells suspended in DMEM or DMEM/F12 was slowly injected into the brain over a 3 minute period (1μL per minute), followed by 3 minutes of rest. Following the injection, the syringe was slowly withdrawn to avoid backflow of cells and the burr hole was filled with bone wax. To visualize GL261-luc tumors, GBM-bearing mice were injected i.p. with 150mg/kg D-luciferin (Fisher Scientific) in 200μL PBS. After 10 minutes, mice were anesthetized with 2% isoflurane and imaged using the IVIS Spectrum In Vivo Imaging System (Xenogen). To establish flank tumors, 1×10^6^ B2905-M4 cells suspended in 50μL of RPMI were injected i.d. into the right flank.

For experiments assessing the contribution of pre-existing T_RM_, depletion of circulating memory OT-I T cells was achieved by injecting 0.25μg of an anti-Thy1.1 depleting Ab (clone HIS51, BD Biosciences 554892) intraperitoneally 1 week prior to tumor injections. Alternatively, FTY720 was injected daily intraperitoneally at a dose of 0.1mg/mL, starting one week prior to tumor injections and continuing for the duration of the experiments thereafter.

For experiments assessing T_CircM_ phenotype following entry into tumors, spleens were harvested from OT-I memory mice and processed as described under *Lymphocyte isolation and phenotyping*. The isolated lymphocytes were enriched for CD8+ T cells using a STEMCELL Technologies EasySep™ Mouse CD8+ T Cell Isolation Kit. An aliquot of isolated CD8+ T cells was then stained with antibodies to determine the frequency of OT-I T cells within the sample. 100,000 circulating memory OT-I T cells were intratumorally (i.t.) injected into GL261, 005, or B2905-M4 tumors or 1,000,000 circulating memory OT-I T cells were i.v. injected into mice 24 hours before tumor injection or into mice already bearing intracranial GL261 tumors. For PTx experiments, enriched CD8+ T cells were cultured in a round-bottom plate with 100ng/ml PTx (Sigma Aldrich) or control media for 60 minutes at 37°C in 5% CO2. Following incubation, cells were washed three times and an aliquot was stained with antibodies to determine the frequency of OT-I T cells within the control and treated samples. The cells were then i.v. injected into mice bearing GL261 tumors, as previously described, and tumors were harvested 24 hours later.

### Luminex assay

Cytokines were measured using Millipore human cytokine multiplex kits (EMD Millipore Corporation, Billerica, MA). Calibration curves from reconstituted cytokine standards were prepared with fourfold dilution steps in the same matrix as the samples. High and low controls from the Millipore kits were included. Standards and controls were measured in triplicate, samples were measured once, and blank values were subtracted from all readings. All assays were carried out directly in a 96-well plate (Millipore, Billerica, MA) at room temperature and protected from light. Briefly, beads together with a standard, sample, control, or blank were added along with their appropriate matrix solution (assay buffer or serum matrix (Millipore, Billerica, MA)) in a final volume of 50μL and incubated together at room temperature for 2 hours with continuous shaking. Beads were washed three times with 200μL 1x Wash Buffer (Millipore, Billerica, MA) using a magnetic plate washer. Detection Antibodies (25μL/well, Millipore, Billerica, MA) were added to the beads for a 1 hour incubation with continuous shaking. No wash after. Then streptavidin-PE (25μL/well, Millipore, Billerica, MA) was added to the beads for an additional 30 minute incubation. Beads were washed three times with 200μL 1x Wash Buffer (Millipore, Billerica, MA) using a magnetic plate washer and resuspended in 150μL of Sheath Fluid (Bio-Rad Cat#171000055) for at least 5 minutes. The fluorescence intensity of the beads was measured using the Bio-Plex array reader. Bio-Plex Manager software with five-parametric-curve fitting was used for data analysis.

### Mouse T cell enrichment and FACS sorting

Lymphocytes were isolated from tumors and spleens as previously described in *Lymphocyte isolation and phenotyping*. 8-10 mice were pooled per tumor condition (T_CircM_-only, T_RM_-only, or Spleen). Each sample was stained for CD8β-StarBright Violet 710 (clone KT15, Bio-Rad 64577111), CD45.1-PE-Cy7 (clone A20, BioLegend 110729), and DAPI, and barcoded with tissue-specific hashtags, TotalSeq™-B0302 anti-mouse Hashtag 2 Antibody (clone M1/42; 30-F11, BioLegend 155833), TotalSeq™-B0303 anti-mouse Hashtag 3 Antibody (clone M1/42; 30-F11, BioLegend 155835), TotalSeq™-B0304 anti-mouse Hashtag 4 Antibody (clone M1/42; 30-F11, BioLegend 155837), before sorting using a Sony Biotechnologies SH-800 Cell Sorter. Viable CD8+ CD45.1+ OT-I T cells, identified by DAPI exclusion, were sorted to >80% purity on a Sony SH-800 Cell Sorter. Following sorting into PBS + 0.5% BSA, cell suspensions were submitted to the Dartmouth Genomics Shared Resource core facility for processing.

### Mouse scRNA-seq library construction and sequencing

Cell suspensions were counted on a Nexcelom K2 automated cell counter with acridine orange/propidium iodide staining. Cells were loaded on a Chromium Single Cell G Chip (10x Genomics Inc.) targeting 10,000 cells/sample. Libraries were prepared using the Chromium Next GEM 3’ (PN-1000269) according to the manufacturer’s protocol (CG000317), producing separate libraries for gene expression and for hashtag-derived barcodes. Libraries were sequenced on an Illumina NextSeq2000 (Illumina, Cat#20040559), using Read1: 28bp, Read2: 90bp. Sequencing depth averaged 60,000 reads per cell for the gene-expression libraries and 10,000 reads/cell for the Hashtag libraries.

### Mouse scRNA-seq data analysis

Raw sequencing reads were demultiplexed and mapped to the mm10-2020-A genome using CellRanger v9.0.0. Standard analysis using Seurat v5.3.1.9999 in R v4.4.3 retained genes expressed in at least three cells and retained cells with fewer than 73,148 total reads/cell, between 500 and 8,520 expressed genes, and less than 4% mitochondrial reads. Multiplexed sample identities were extracted from CLR-normalized Hashtag reads using a 99% positive quantile. Additional T_RM_-only tumor cells were recovered among cells with non-Singlet calls when sample_maxID=’ T_RM_ Only’ by imposing manual thresholds of greater than four reads among the Doublet calls and greater than one read among the Negative calls.

RNA reads were normalized with SCTransform, regressing on the percent mitochondrial reads. TCR genes, Gm42418, AY036118, and Lars2 were removed from among the variable features. The Uniform Manifold Approximation and Projection (UMAP) and Louvain clustering with 0.6 resolution were performed on 30 principal components. Small clusters of contaminating cells were removed to yield eight clusters among 4,677 cells (1,540 T_RM_-only tumor, 1,400 T_CircM_-only tumor, 1,737 Spleen). Cell cluster identities were determined by manual inspection of top marker genes. Volcano plots were computed with SCTransform data using the SeuratExtend v1.2.4 R library. Features plots, violin plots, dot plots and module scoring utilized data imputed using the ALRA v0.0.0.9000 R library. Residency and Circulating module scores were computed using published human gene signatures (22,39) (specifically Table S2 Non-exclusive TRMup/dn lists) converted to mouse genes using the homologene v1.4.68.19.3.27 R library.

### Human blood and tissue procurement, processing, and FACS sorting

The GBM tumor sample, peritumoral brain, and paired blood were obtained from a standard-of-care surgical resection, with tumor and peritumoral brain tissue confirmed by the Department of Pathology at Dartmouth Health (Lebanon, NH) according to the Institutional Review Board guidelines. Blood samples were diluted 1:1 in PBS and underlaid with 10 ml of Ficoll (Cytiva) in a 50ml conical tube, then spun for 30 min at 550 × g at room temperature. The PBMC layer was collected into a 50ml conical tube, diluted with RPMI 1640, and spun at 690 × g for 10 min. Tumor and peritumoral brain were cut into small pieces and enzymatically dissociated with 0.5mg/ml Collagenase IV (Sigma) and DNAse I (5mg/ml) for 30 minutes at 37°C. Tissues were then mechanically disrupted using a GentleMacs dissociator (Miltenyi Biotec), filtered through 70μm mesh, and isolated using a 44/67% Percoll (Cytiva) density gradient. The enriched lymphocyte fraction was then collected and washed prior to staining with CD4-BV711 (clone RPA-T4, BioLegend 300558), CD8-PE-Cy7 (clone SK1, BioLegend 344712), and barcoded with tissue-specific hashtags, TotalSeq™-C0251 anti-human Hashtag 1 Antibody (LNH-94; 2M2, BioLegend 394661), TotalSeq™-C0252 anti-human Hashtag 2 Antibody (LNH-94; 2M2, BioLegend 394663), and TotalSeq™-C0253 anti-human Hashtag 3 Antibody (LNH-94; 2M2, BioLegend 394665) for 30 minutes at 4°C in PBS + 2% FBS. Viable CD8+ cells, identified by DAPI exclusion, were sorted to >80% purity on a Sony SH-800 Cell Sorter. Following sorting into PBS + 0.5% BSA, cell suspensions were submitted to the Dartmouth Genomics Shared Resource for processing.

### Human TCR-seq data analysis

Raw sequencing reads were demultiplexed and mapped using the GRCh38-2024-A and vdj-GRCh38-alts-ensembl-5.0.0 references with CellRanger v9.0.0. Standard analysis of the transcriptomic data using Seurat v5.3.1.9999 in R v4.4.3 retained genes expressed in at least three cells and retained cells with fewer than 10,000 total reads/cell, between 500 and 4000 expressed genes, and less than 10% mitochondrial reads. Multiplexed sample identities were extracted from CLR-normalized Hashtag reads using a 95% positive quantile. RNA reads were normalized with SCTransform, regressing on the percent mitochondrial reads.

The VDJ data were analyzed using scRepertoire v2.7.3 in R v4.5.3 to combine contig lists with parameters set to not filter out NA or multi chains, yet to filter out nonproductive chains. Clones shared among tissues were identified using clonalCompare(), where a clone was defined as the nucleotide sequence of the TCR beta CDR3 region. Venn diagrams were computing using ggVennDiagram v1.5.7. Differential gene expression in cells for a given subset of clones and tissues was assessed using the Wilcoxon test in the Seurat FindMarkers() function.

### Cell definitions

Definition of T cell nomenclature used in this paper is outlined here and was determined by either flow cytometry phenotyping or by corresponding transcriptional signatures in sequencing data (53). T cells were considered naïve if they had not yet experienced stimulation with cognate antigen, or if CD44^low^ following antigen experience. T cells were classified as ‘activated’ or ‘effectors’ if recently stimulated with cognate antigen (<14 days post-acute infection), and as ‘resting memory’ if CD44^hi^ >30 days post-acute infection. T_CM_ (also referred to as CD8+ TSM) were defined as CD44^hi^/CD62L+/CD69-/CD103-isolated from blood, spleen, or tumor, or transcriptionally by elevated expression of *Ccr7* and *Sell*. T_EM_ (also referred to as TD_B_M) were defined as CD44^hi^/CD62L-/CD69-/CD103-cells isolated from blood or spleen, or transcriptionally by low expression of *Ccr7* and *Sell*, and low expression of residency-associated genes outlined in Fig. 6. T_CircM_ were defined as all CD44^hi^/CD69-/CD103-memory T cells isolated from blood or spleen. T_RM_ (also referred to as TD_R_M) were defined as i.v.-label-negative cells expressing CD69 and/or CD103 isolated from brain or skin based on previously published parabiosis experiments (33,54). ‘T_RM_-like’ cells were defined as CD8+ T cells from tumors expressing CD69+ CD103+/-(by protein) or with transcriptional signatures of T_RM_. They have not been assessed for migratory properties and therefore true tissue residence isn’t claimed. The term ‘bystander’ refers to virus-specific T cells that infiltrated wild-type tumors and therefore did not encounter cognate antigen.

### Statistical analysis

Statistical analysis was performed using GraphPad Prism software. Details of biological replicates, experimental replicates, and statistical tests used for each dataset are provided in the figure legends. Normality of each dataset was assessed with the Shapiro-Wilk test. For data that passed the normality test, paired *t* tests were applied to paired samples and unpaired *t* tests or Welch’s t test (unequal variance) were performed on unpaired samples. When normality was not met, Mann-Whitney test was performed. For experiments involving more than two groups, one-way or two-way ANOVA with a Tukey posttest was used for normally distributed data and Kruskal-Wallis test with Dunn’s post-hoc analysis was used for nonparametric data. All results are expressed as mean ± SEM, and statistical significance was set at *p* < 0.05.

## Acknowledgements

We thank the members of the Rosato, Skorput, and Turk labs at Dartmouth for valuable discussions and collaboration. We are grateful to Aaron Christensen for graphical assistance. VSV_OVA_ was generously provided by Dr. David Masopust. GL261-luc cells were generously shared by Dr. Tyler Curiel. B2905-M4 cells were generously provided by Dr. Mary Jo Turk. 005 cells were generously gifted by Dr. Susan Kaech. We thank the clinical coordinators at Dartmouth Health for their efforts in consenting patients and facilitating the collection of tissue specimens from routine surgical resections. We are deeply grateful to the patients and their families who generously agreed to participate in this study and contribute tissue samples. The authors acknowledge the use of shared resources at the Dartmouth Cancer Center, supported by the NCI Cancer Center Support Grant P30CA023108. Specifically, we utilized the Genomics and Molecular Biology Shared Resource (RRID:SCR_021293), including Genomics and 10x Chromium Single Cell & Spatial Genomics services, supported by grants S10OD030242, P20GM130454, and S10OD025235. We also utilized the Immune Monitoring and Flow Cytometry Shared Resource (RRID:SCR_019165). These shared resources are housed within the Dartmouth Cancer Center at Dartmouth College. We acknowledge the DartLab Immune Monitoring Core and the Genomics and Molecular Biology Shared Resource core facility for maintaining instruments and for their technical assistance, as well as Dartmouth Animal Care Facility staff members for animal husbandry. Studies were supported through funding from the Dartmouth Cancer Center and the National Institutes of Health (NIH) grants R01AG078761 and R01CA269455 to PCR, R01CA22502 to MJT, R01CA254042 to MJT and YHH, Friends of Dartmouth Cancer Center funding (The Nelson Andrea Clark Medical Research fund, The Anderson Charles H Cancer Research Fund, and ICI COMPEL award) to P.C.R and S.S.G. Generative AI was used to assist with wording and clarity.

## Author contributions

Conceptualization: SAK, PCR Methodology: SAK, PCR Analysis: SAK, SML

Investigation: SAK, TC, SCM, NRD, HND, SCB, MAF, JFI, ACR, AJS

Visualization: SAK, PCR Funding acquisition: SSG, PCR Project administration: PCR

Resources: CVA, CCL, NES, LTE, SSG, MJT, AGJS, PCR

Supervision: PCR

Writing – original draft: SAK, PCR

Writing – review & editing: SAK, TC, SCM, SSG, MJT, PCR

## Data and materials availability

All materials, including cell lines and reagents, will be made available upon request. No original code was created in this manuscript. All packages and scripts were modified from publicly available sources for case-specific uses. All data and code are available upon request. All sequencing datasets generated for this study will be deposited at GEO.

