## Supplemental Figures 1-5 for "Circulating and brain-resident memory CD8+ T cells seed distinct bystander T_RM_-like populations in glioblastoma"

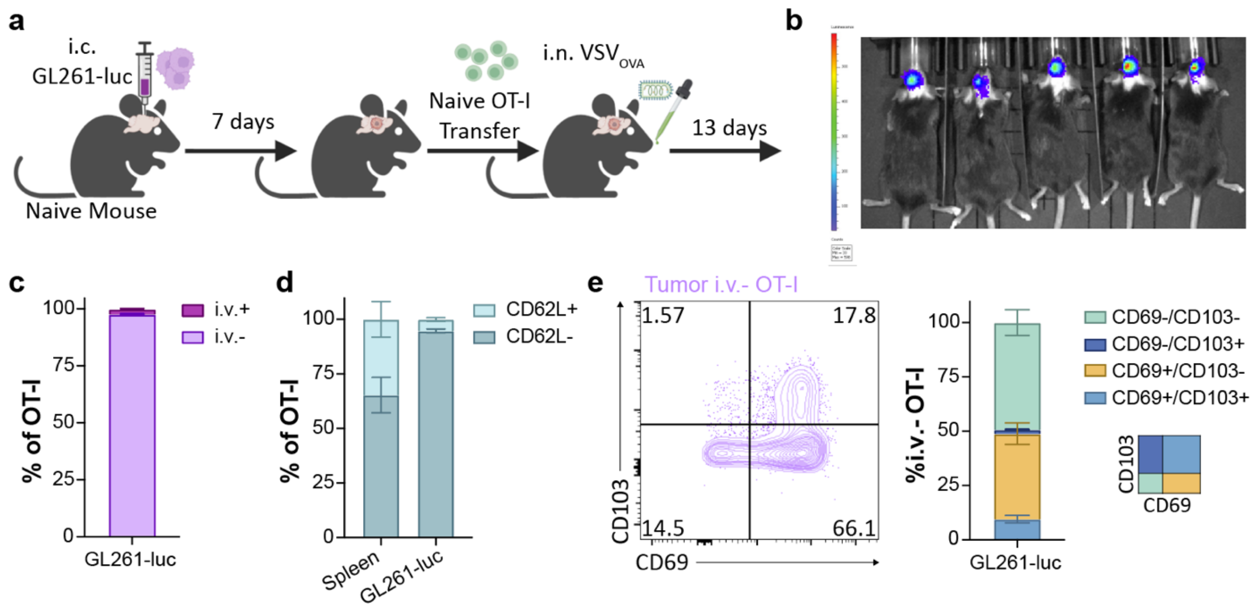


**Supplemental Figure 1. Intranasal infection primes CD8+ T cells to become resident-like in established brain tumors. a)** Schematic of intracranial GL261-luc injection followed by OT-I T cell transfer and intranasal infection with $VSV_{OVA}$. **b)** Representative IVIS bioluminescence images of GL261-luc-bearing mice 7 days post-i.c. injection. **c)** Bar graph depicting i.v. antibody label of OT-I T cells isolated from GL261-luc tumors. **d)** Bar graphs showing distribution of CD62L- and CD62L+ $T_{CircM}$ isolated from the spleen and tumor. **e)** Representative flow plot and quantification of CD69 and CD103 expression on tumor-infiltrating OT-I T cells 13 days post-intranasal infection (20 days post-tumor challenge). Data in b) – e) are combined from two independent experiments; $n = 9$. Error bars represent the mean ±SEM.

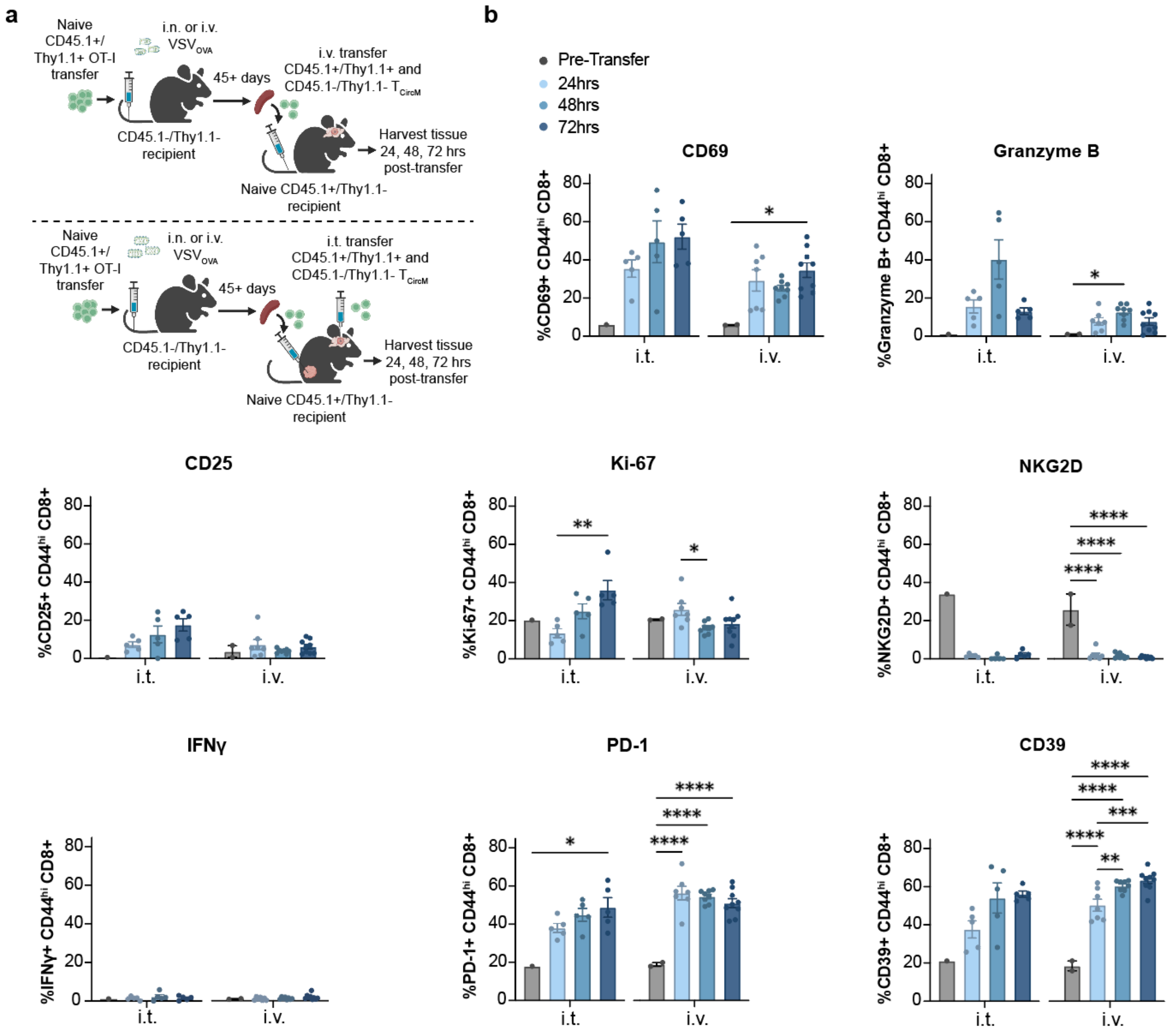


**Supplemental Figure 2. The tumor microenvironment rapidly alters the phenotypes of antigen-experienced endogenous CD8+ T cells. a)** Schematic of i.t. or i.v. transfer of $T_{CircM}$ to GL261 tumor-bearing mice. **b)** Expression of markers associated with residency and cytokine-mediated activation on CD44$^{hi}$ CD8+ T cells before (spleen) and 24, 48, and 72 hours after i.v. transfer (tumor). I.t. data in (b) are from one experiment; Pre-transfer, $n = 1$; 24hrs, $n = 5$; 48hrs, $n = 5$; 72hrs, $n = 5$. I.v. data in (b) are combined from two independent experiments; Pre-transfer, $n = 2$; 24hrs, $n = 7$; 48hrs, $n = 8$; 72hrs, $n = 9$. Error bars represent the mean ±SEM with $p$ values determined by one-way ANOVA or Kruskal-Wallis test for non-normally distributed data. *$p < 0.05$, **$p < 0.01$, ***$p < 0.001$, ****$p < 0.0001$.

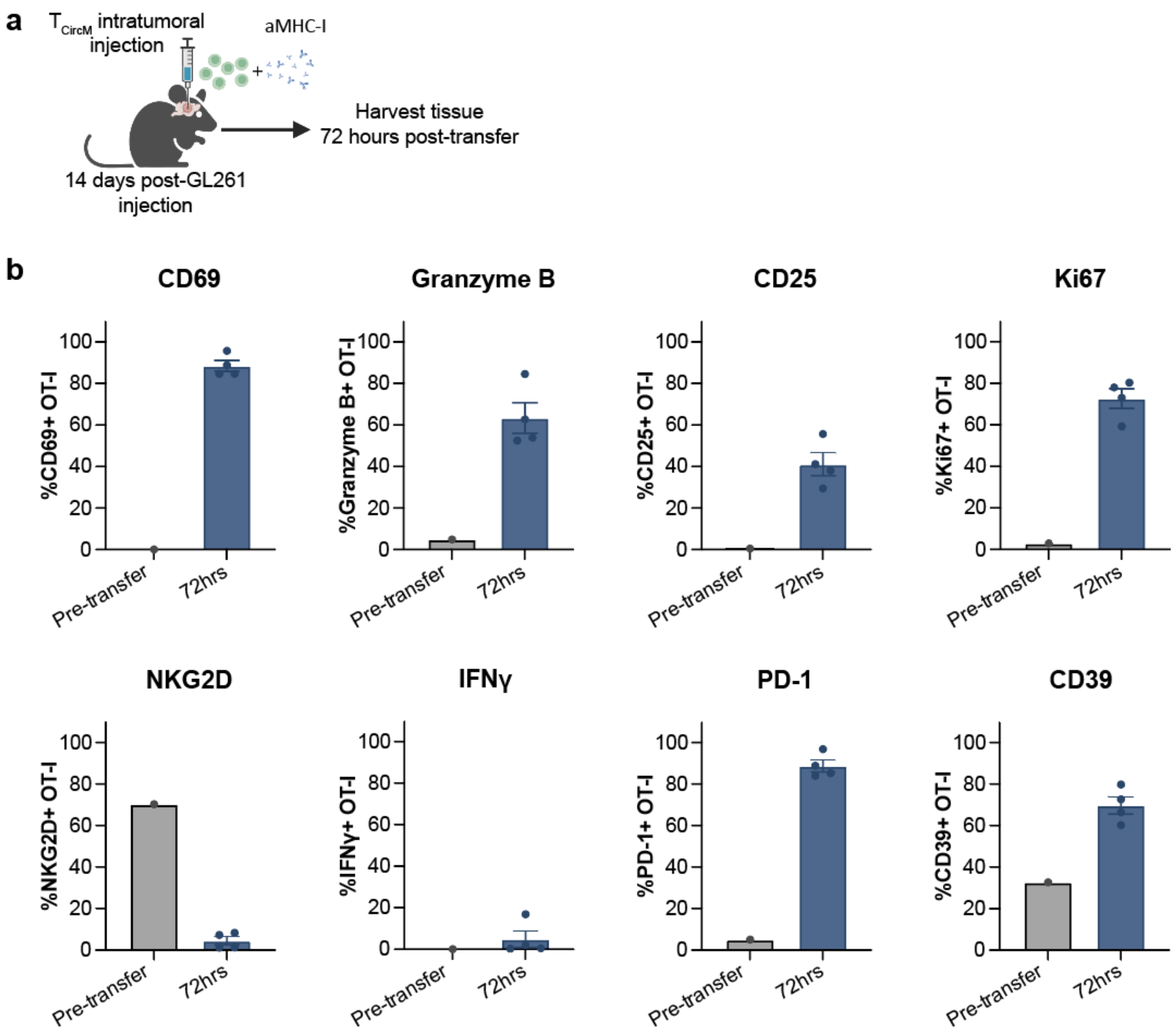


**Supplemental Figure 3. The altered phenotype of $T_{CircM}$ upon tumor entry is sustained in the presence of MHC-I blocking antibody. a)** Schematic of i.t. transfer of $T_{CircM}$ + αMHC-I antibody to GL261 tumor-bearing mice. **b)** Frequency of OT-I expressing markers associated with residency and cytokine-mediated activation out of total OT-I T cells before (spleen) and 72 hours after i.t. transfer (tumor). Data are from one experiment; Pre-transfer, $n = 1$; 72hrs, $n = 4$.

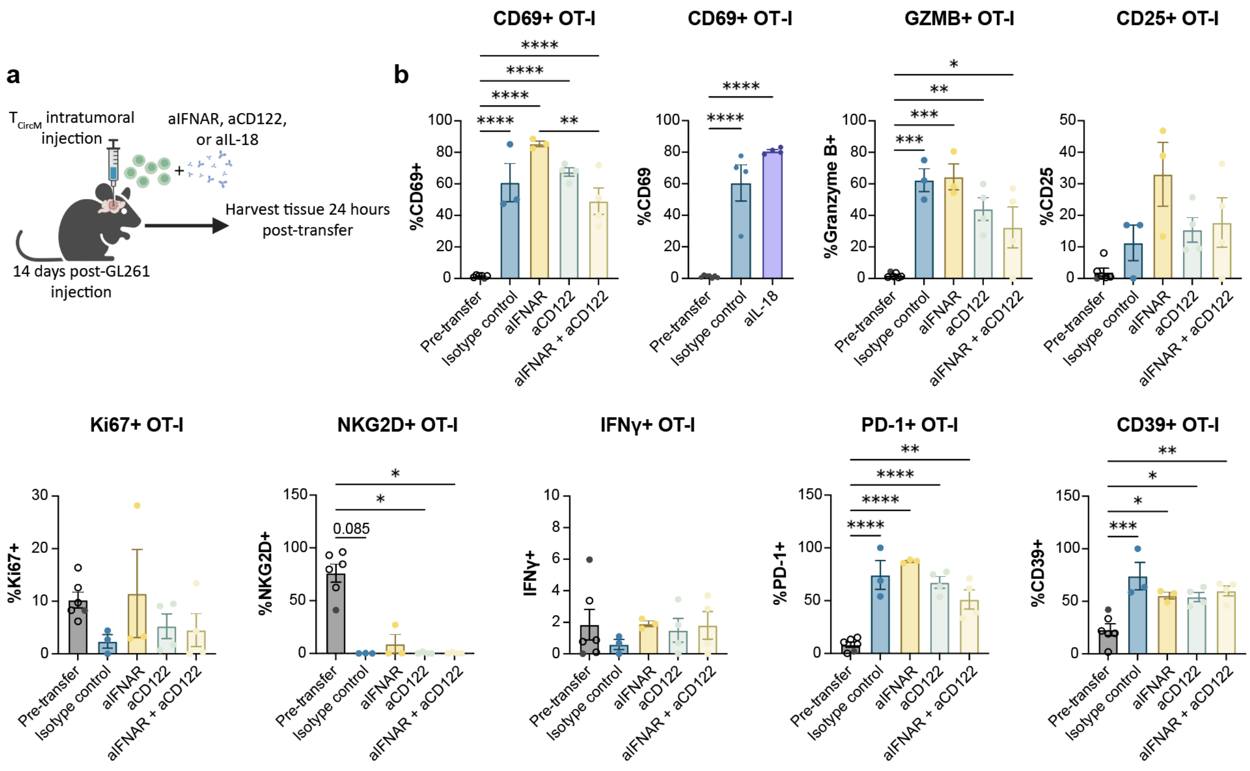


**Supplemental Figure 4. The altered phenotype of $T_{CircM}$ upon tumor entry is sustained in the presence of cytokine blockade. a)** Schematic of i.t. transfer of $T_{CircM}$ incubated with αIFNAR, αCD122, or αIL-18. **b)** Frequency of OT-I T cells expressing markers associated with residency or cytokine-mediated activation out of total OT-I T cells before (spleen) and 24 hours after i.t. transfer (tumor). Data are from one experiment; Pre-transfer, $n$ = 6; Isotype control, $n$ = 3-4; αIFNAR, $n$ = 3; αCD122, $n$ = 4; αIFNAR + αCD122, $n$ = 4; αIL-18, $n$ = 4. Pre-transfer phenotypes of OT-I splenocytes (pooled from all experiments) were obtained from unmanipulated OT-I memory mouse splenocytes. Closed dots indicate cells used in the experiment shown; open circles represent cells from experiments with a different endpoint. Error bars represent the mean ±SEM with $p$ values determined by one-way ANOVA or Kruskal-Wallis test for non-normally distributed data. *$p < 0.05$, **$p < 0.01$, ***$p < 0.001$, ****$p < 0.0001$.

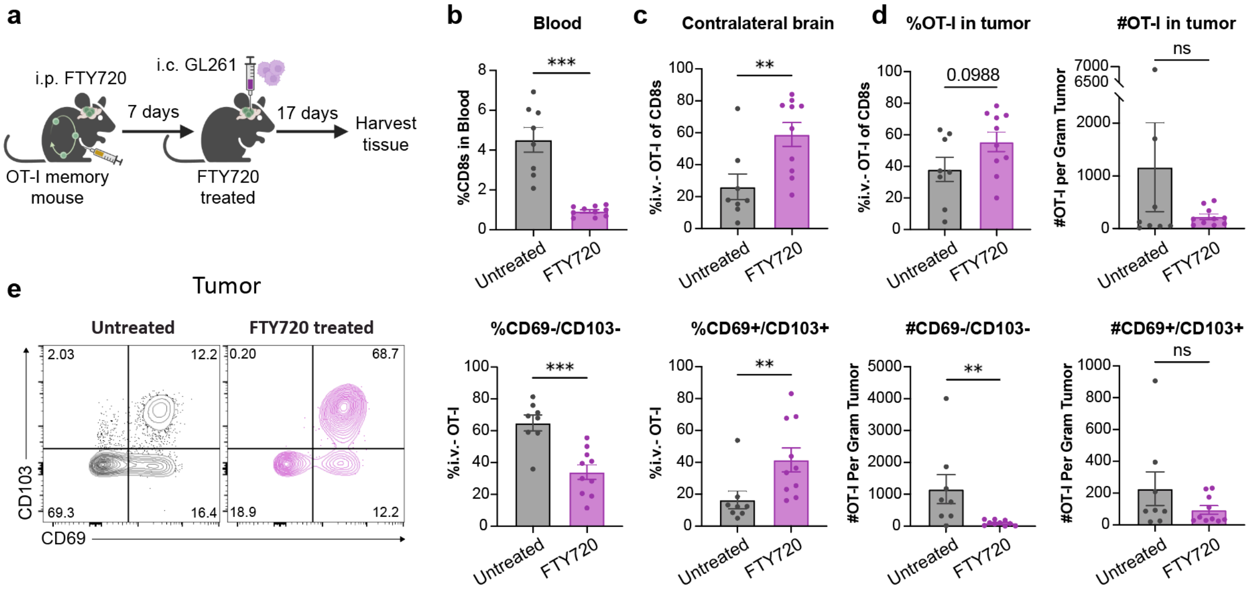


**Supplemental Figure 5. The GBM CD103+ TIL compartment is enriched in the presence FTY720. a)** Schematic of FTY720 treatment and tumor implantation. **b)** Frequency of CD8+ T cells in the blood following FTY720 treatment, prior to tumor injection. **c)** Frequency of i.v.-negative OT-I T cells isolated from the contralateral brain tissue. **d)** Frequency and number of OT-I T cells isolated from tumors. **e)** Representative flow plots and quantification of i.v.-negative tumor-infiltrating OT-I T cells in untreated and FTY720-treated mice. Data are combined from two independent experiments; Untreated, $n = 8$; FTY720, $n = 10$. Error bars represent the mean ±SEM with $p$ values determined by unpaired $t$ test or Mann Whitney test for non-normally distributed data. **$p < 0.01$, ***$p < 0.001$. ns, not significant.
